# A novel computational method to design BH3-mimetic peptide inhibitors that can bind specifically to Mcl-1 or Bcl-X_L_

**DOI:** 10.1101/2020.07.09.194662

**Authors:** C. Narendra Reddy, Nishat Manzar, Bushra Ateeq, Ramasubbu Sankararamakrishnan

## Abstract

Interactions between pro- and anti-apoptotic B cell lymphoma 2 (Bcl-2) proteins decide the fate of the cell. BH3 (Bcl-2 homology 3) domain of pro-apoptotic Bcl-2 proteins interacts with the exposed hydrophobic groove of anti-apoptotic counterparts. Design and development of BH3 mimetics that target the hydrophobic groove of specific anti-apoptotic Bcl-2 proteins have the potential to become anti-cancer drugs. We have developed a novel computational method to design sequences with BH3 domain features that can bind specifically to anti-apoptotic Mcl-1 or Bcl-X_L_. In this method, we retained the four highly conserved hydrophobic and aspartic residues of wild-type BH3 sequences and randomly substituted all other positions to generate a large number of BH3-like sequences. We modeled 20000 complex structures with Mcl-1 or Bcl-X_L_ using the BH3-like sequences derived from five wild-type pro-apoptotic BH3 peptides. Peptide-protein interaction energies calculated from these models for each set of BH3-like sequences resulted in negatively-skewed extreme value distributions. The selected BH3-like sequences from the extreme negative tail regions have distinctly different distribution of charged residues for Mcl-1 and Bcl-X_L_. BH3-like sequences with highly favorable interaction energies prefer to have acidic residues for Mcl-1 and are enriched with basic residues when they bind to Bcl-X_L_. With the charged residues often away from the binding interface, the overall electric field generated by the charged residues result in highly favorable long-range electrostatic interaction energies between the peptide and the protein giving rise to high specificity. Cell viability studies of representative BH3-like peptides further validated the predicted specificity.

## Introduction

Intrinsic pathway of apoptosis is regulated by more than 20 proteins belonging to the B cell lymphoma 2 (Bcl-2) family ^1-3^. Balance between the cell survival and death is maintained by the expression levels of different classes of Bcl-2 members ^4^. Bcl-2 family can be largely classified into three groups depending upon the number of conserved Bcl-2 homology (BH) domains. Anti-apoptotic Bcl-2 proteins (Bcl-X_L_, Bcl-2, Bcl-W, Mcl-1 and A1) have four BH regions (BH1 to BH4). The mitochondrial pore-forming BAX and BAK with multi-BH domains and the “BH3-Only” activator proteins such as BIM, BID and PUMA are pro-apoptotic and constitute two other classes of the Bcl-2 family ^5^. The pro-apoptotic activator proteins trigger apoptosis by binding with the anti-apoptotic Bcl-2 members and also activate BAX and BAK leading to oligomerization ^6^. These oligomers eventually form pores in the mitochondrial outer membrane. This process called mitochondrial outer membrane permeabilization (MOMP) results in the release of mitochondrial proteins such as cytochrome C activating the downstream of intrinsic apoptotic pathway ^7^. Overexpression of anti-apoptotic Bcl-2 members and mutations that compromised the function of pro-apoptotic members have been reported in several human cancers making them as attractive targets for anti-cancer drugs ^8-9^. Since the interactions between anti- and pro-apoptotic Bcl-2 members are vital for the cells to die or survive, understanding the protein-protein interactions among the Bcl-2 family members have assumed prime importance among the researchers. The most important step in this direction is to determine the three-dimensional structures of anti-apoptotic Bcl-2 proteins in complex with the pro-apoptotic counterparts. Structures of the anti-apoptotic Bcl-2 family members have been determined either in the apo-form or in complex with pro-apoptotic BH3 peptides or small molecules ^10-11^. Anti-apoptotic Bcl-2 members exhibit a conserved helical fold despite the low sequence identity among themselves ^10, 12^. They are characterized by seven to eight α-helices with the central hydrophobic helix surrounded by other helices which are mostly amphipathic in nature. The characteristic feature of Bcl-2 fold is the pronounced hydrophobic groove formed by the BH1 to BH3 domains. BH3 region forms amphipathic α-helix and the hydrophobic part of the BH3 helical region of pro-apoptotic Bcl-2 members binds to this hydrophobic groove of anti-apoptotic Bcl-2 proteins. Experiments have revealed that the pro-apoptotic BH3 domain is capable of eliciting the desired biological response of the full protein ^13-14^. Hence many structures included the anti-apoptotic Bcl-2 members in complex with BH3 peptides derived from the pro-apoptotic members.

Resistance to apoptosis due to intrinsic pathway is mediated by anti-apoptotic Bcl-2 proteins and hence targeting them is one of the important strategies to find therapeutic solutions for different cancers. Some of the strategies including NMR-based screening and structure-based drug design are being pursued to find molecules that selectively target specific anti-apoptotic proteins ^2, 15^. Inhibitors could be small molecules or peptides mimicking the BH3 region of pro-apoptotic Bcl-2 proteins which can activate the intrinsic apoptotic pathway by interacting with the anti-apoptotic Bcl-2 proteins. These are collectively called BH3-mimetics and a significant amount of research has been carried out to develop BH3-mimitetics. Early breakthroughs in this direction include discovery of inhibitors such as ABT-737 ^15^, navitoclax (ABT-263) ^16^ and venetoclax (ABT-199) ^17^. The developed inhibitors fall into mainly two classes. The first class of molecules targets Bcl-X_L_, Bcl-2 or at least one of the three anti-apoptotic proteins, Bcl-2, Bcl-X_L_ and Bcl-W. The second class of inhibitors such as UMI-77 ^18^ and S63845 ^19^ target only Mcl-1. Several of these molecules that inhibit Bcl-X_L_ or Bcl-2 have entered pre-clinical and clinical trials. For example, the Bcl-2-specific venetoclax is already in Phase-I or Phase-II trails either as a single agent or in combination with other drugs ^20-21^. The use of venetoclax in chronic lymphocytic leukaemia has been recently approved by the US Food and Drug Administration ^20,22^.

The binding of BH3 region of pro-apoptotic Bcl-2 proteins to the hydrophobic groove of anti-apoptotic Bcl-2 partners is possible due to the four conserved hydrophobic residues in the hydrophobic side of BH3 amphipathic helix. Electrostatic interactions due to a conserved Asp provide additional stabilization for the protein-protein complex structure. Amy Keating and her group used yeast surface display and SPOT peptide arrays to isolate and characterize BH3 peptide mutants that bind specifically to Mcl-1 or Bcl-X_L_ ^23^. They identified specific positions that are responsible for binding to Mcl-1 or Bcl-X_L_. The conclusions of this study were mainly based on the SPOT experiments and not based on a rigorous systematic binding affinity studies. Stewart et al. tested BH3 peptides derived from different pro- and anti-apoptotic Bcl-2 proteins stabilized by hydrophobic staples. They showed that the BH3 helix derived from Mcl-1 is the only peptide among the tested peptides that showed high selectivity for Mcl-1 ^24^. They could also explain the selectivity by specific interactions between the highly conserved BH3 positions and the Mcl-1 protein. Foight et al. engineered three highly Mcl-1-specific Bim-based BH3 peptides and demonstrated their usefulness in BH3 profiling assays ^25^. A hydrophobic staple scan helped to optimize the staple positions that conferred selective binding to Mcl-1 with high affinity ^26^. These peptide constructs provided resistance to protease activity and were shown to be cell-penetrant. Structural characterization of the designed peptides in complex with the specific anti-apoptotic proteins were carried out in many of these studies ^24,26-29^.

Although the above studies have been successful to some extent, it is still difficult to rationally design BH3-like peptides that can bind to a specific anti-apoptotic Bcl-2 protein with high affinity. In this study, we have developed a computational method to identify BH3-like peptides from a large number of randomized BH3-like sequences that can bind with high affinity either with Bcl-X_L_ or Mcl-1. We have validated our approach with cell proliferation and cell viability experiments. We also performed several independent molecular dynamics simulation studies on selected peptides for a cumulative period of 10 μs to study the protein-peptide interactions. Our studies show that peptides derived from pro-apoptotic wild-type BH3 sequences when binding to Mcl-1 have distinct preference for acidic residues while those that bind to Bcl-X_L_ are enriched with basic residues. For high-binding affinity, long-range electrostatic interactions between the peptide residues and the protein have important role to play and these residues need not be located in the protein-peptide binding interface. It is surprising since most of the studies focused on substituting amino acids at the binding interface and overlooked the fact that even residues from the solvent-exposed hydrophilic side of amphipathic BH3 helix can play a significant role in giving rise to high affinity for a specific anti-apoptotic protein.

## Results

### Deriving BH3-like sequences and building their complex structures

We first generated BH3-like sequences based on the wild-type BH3 regions of five pro-apoptotic proteins, Puma, Noxa, Bad, Bid and Bim. Two sets (Set-I and Set-II) of one thousand BH3-like sequences were generated for each of the five pro-apoptotic BH3 peptide. When BH3-like sequences were generated, the residues at the four conserved hydrophobic positions and the conserved Asp residue were retained (Figure 1). Residues corresponding to all other positions were randomly chosen. These sequences were used to model the Mcl-1 and Bcl-X_L_ complex structures. With two sets of 1000 sequences generated for each of the five wild-type BH3 peptides and two proteins, a total of 20,000 modeled complex structures built. The modeled structures were used to calculate the interaction energies between the BH3-like peptide and the proteins Mcl-1 or Bcl-X_L_ as described in the Methods section. Histograms of interaction energies were plotted for each library for Mcl-1 and Bcl-X_L_ complexes and are shown in Figure 2 for Set-I. The same data for the second set of BH3-like sequence libraries are presented in the Supplementary Figures for Mcl-1 and Bcl-X_L_ complexes (Figure S1). It is clear that the histograms of interaction energies for each library of BH3-like sequences are not normal distributions and are negatively skewed. These distributions can be described by Gumbel extreme value distribution. This can be observed for both libraries (Set-I and Set-II) of BH3-like sequences generated from the BH3 regions of five pro-apoptotic proteins and both Mcl-1 and Bcl-X_L_ complexes display the same behavior. In each of the histogram, the mean value and the interaction energy obtained for the wild-type peptide are indicated by arrows. With the exception of Mcl-1:Bim and Bcl-X_L_:Bad complexes, interaction energies of all the wild-type peptides are either higher than (less favorable) the mean value or within one standard deviation from the mean value.

**Figure 1:**
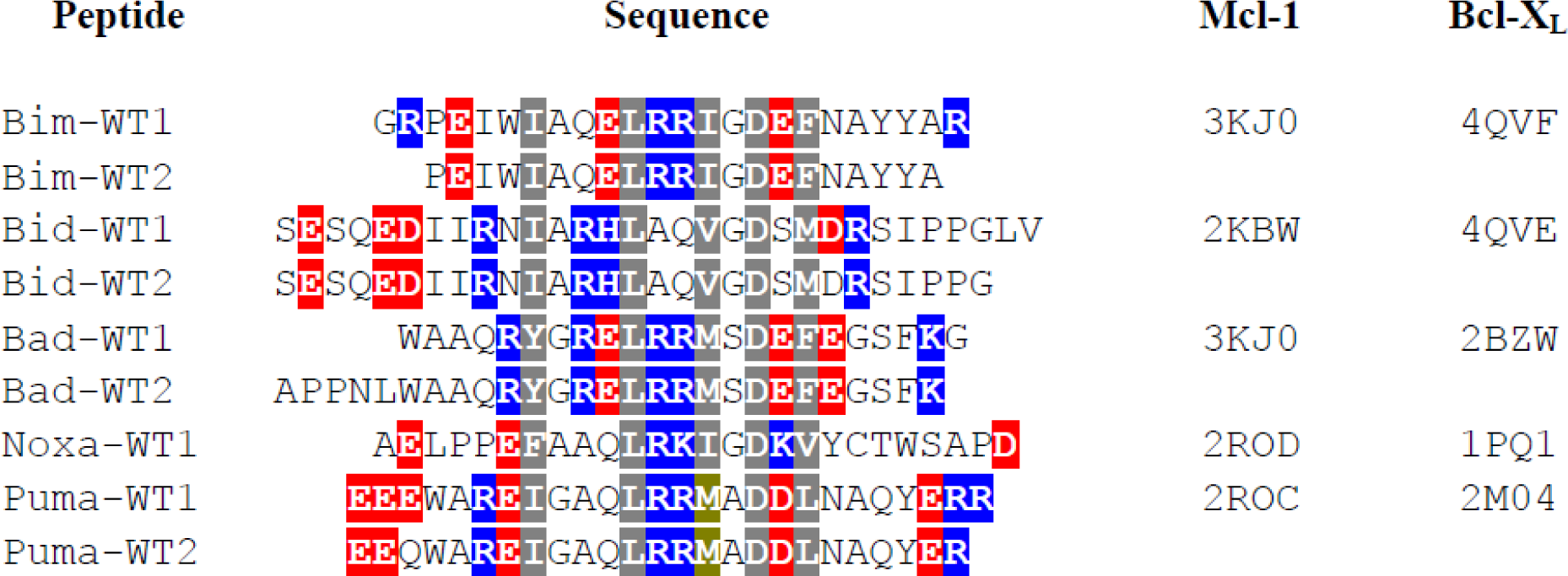
Sequences of wild-type BH3 peptides from the pro-apoptotic proteins Bim, Bid, Bad, Noxa and Puma. The four conserved hydrophobic residues and the conserved Asp are shown in gray background. Acidic and basic residues are displayed in red and blue backgrounds respectively. PDB IDs of the template structures used to model Mcl-1 and Bcl-X_L_ proteins in complex with BH3-like peptide sequences are provided. Reasons for considering two sequences which differ only in length for the BH3 wild-type peptides are explained in the text.

**Figure 2:**
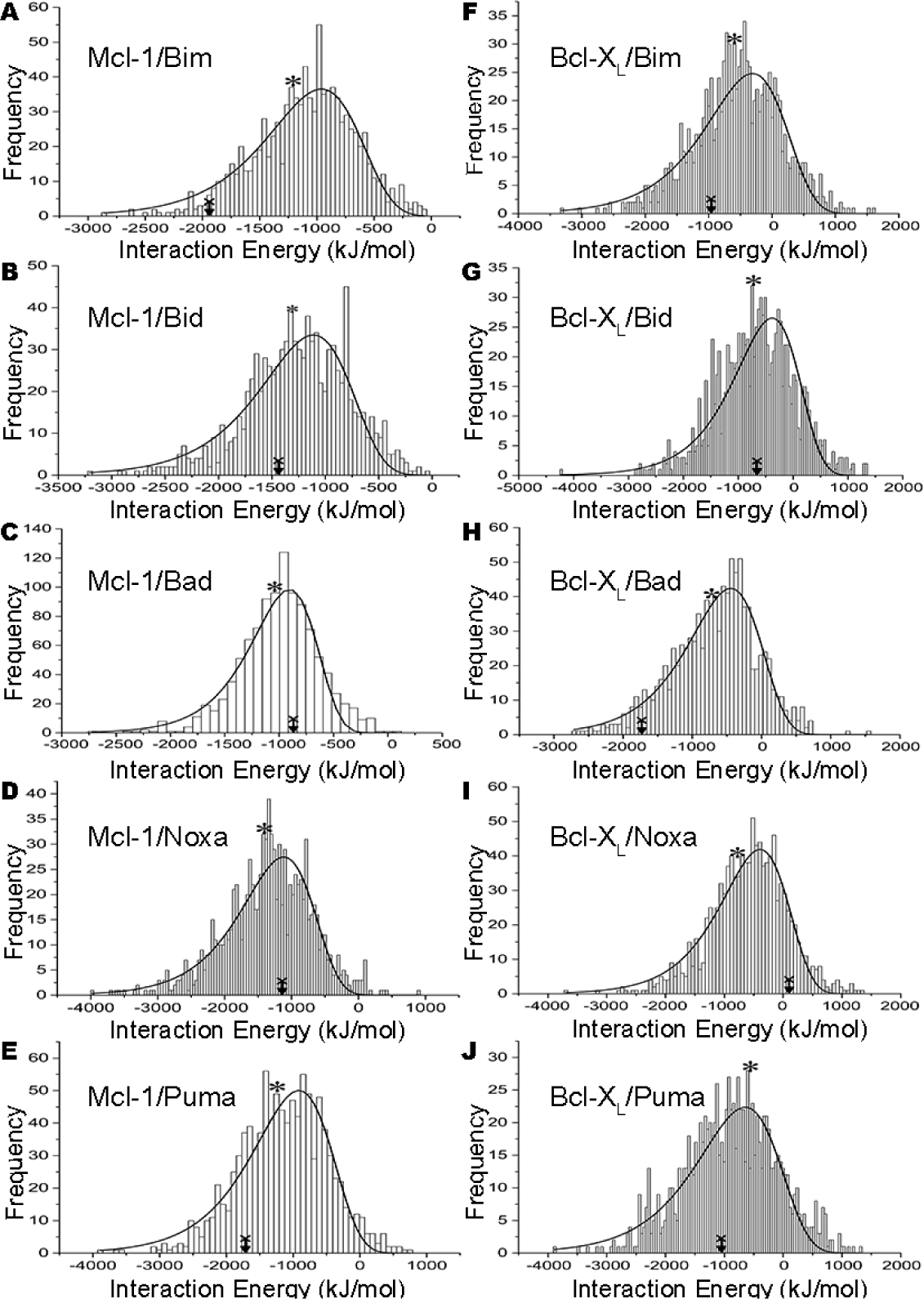
Histograms of interaction energies calculated between BH3-like peptides in complex with (A-E) Mcl-1 and (F-J) Bcl-X_L_. The data shown here correspond to the Set-I of BH3-like peptide sequences derived from the wild-type peptides (A, F) Bim, (B, G) Bid, (C, H) Bad, (D, I) Noxa and (E, J) Puma. The mean value and the interaction energy of the wild-type BH3 peptide are marked in each histogram.

Our interest is to consider the BH3-like sequences whose interaction energies fall within the extreme negative tail of the distribution. Interaction energies of these peptides fall outside two to three standard deviation of the mean value. Probability of finding BH3-like sequences with most favorable interaction energies falling in this region by chance by any random sequence will be extremely low. Hence, we selected the top 50 BH3-like peptides from this region and found out the helix-forming tendency and the hydrophobic moment for each of these peptides as described in the Methods section. For each category of BH3-like peptides derived from a specific pro-apoptotic BH3 region, the top five BH3-like peptides with the highest helical propensity were considered and the sequences are shown in Figure 3 and Figure 4 for Mcl-1 and Bcl-X_L_ complexes respectively. The top Mcl-1-binding BH3-like sequences derived from the Bim wild-type BH3 peptides selected from the first and second sets based on helical propensities are labeled as Bim1-1 to Bim1-5 and Bim2-1 to Bim2-5 respectively. Similarly, the top Bim-based BH3-like sequences with high helical propensities and more favorable interaction energies with Bcl-X_L_ from the first and second sets are labeled as Bim3-1 to Bim3-5 and Bim4-1 to Bim4-5 respectively. A similar convention has been used for BH3-like sequences derived from other wild-type BH3 sequences also. Histograms showing the helix-forming tendencies and helix hydrophobic moments of the selected top BH3-like peptides are plotted along with the wild-type BH3 peptides in the Supplementary Figure S2-S3 and Figure S4-S5 respectively for both sets.

**Figure 3:**
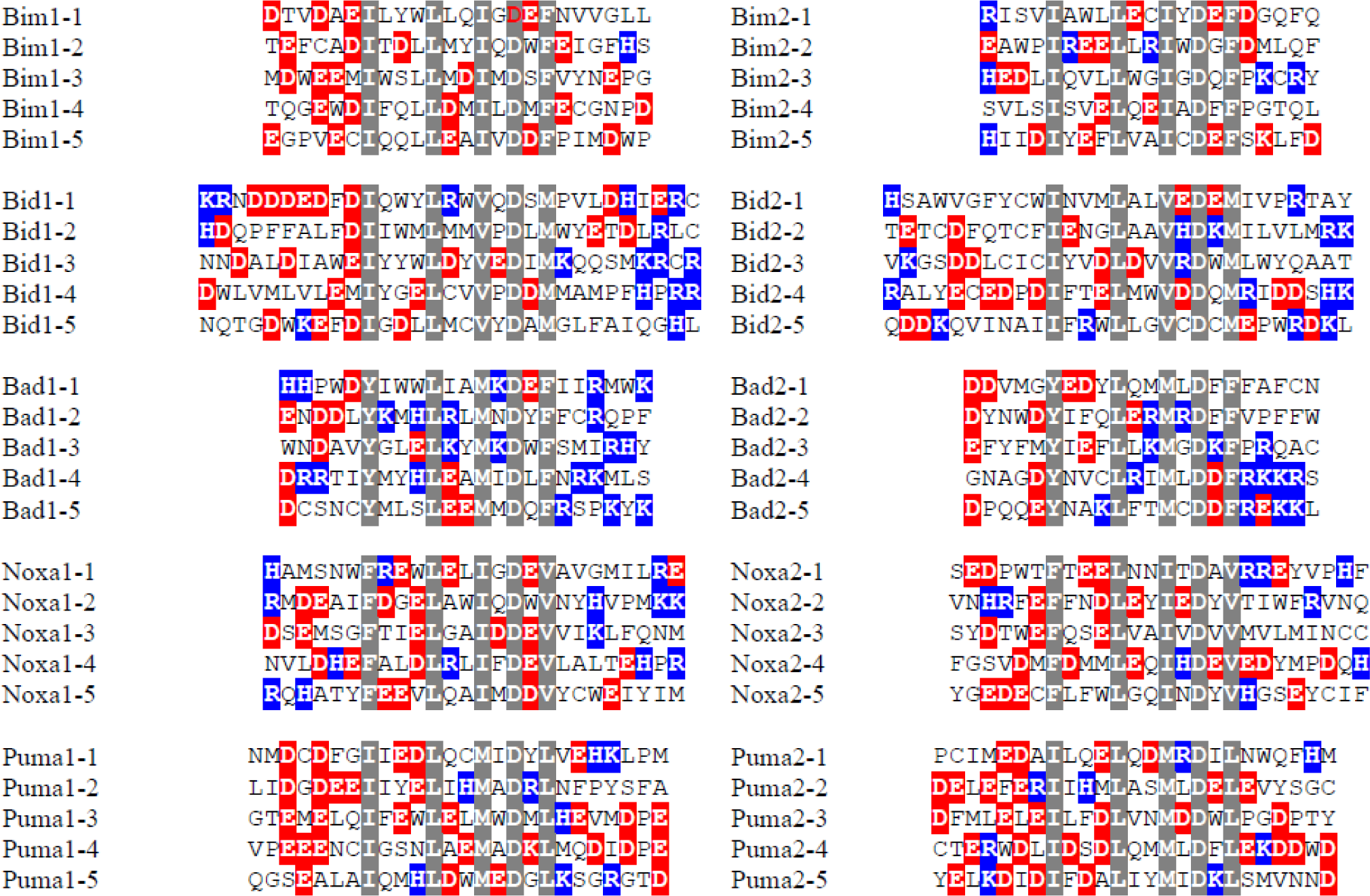
Top BH3-like sequences derived from the wild-type BH3 peptides with highly favorable interaction energies with Mcl-1. Interaction energies of these peptides lie at the negative tail end of the histograms shown in Figure 2 and Figure S1. Those peptides with higher helical propensities and larger hydrophobic moments are selected among the BH3-like sequences that have the most favorable interaction energies with Mcl-1. The four conserved hydrophobic residues and the conserved Asp residue are shown in gray background. The acidic and basic residues are shown in red and blue backgrounds respectively. The sequences shown on the left and right sides correspond to those from Set-I and Set-II respectively.

**Figure 4:**
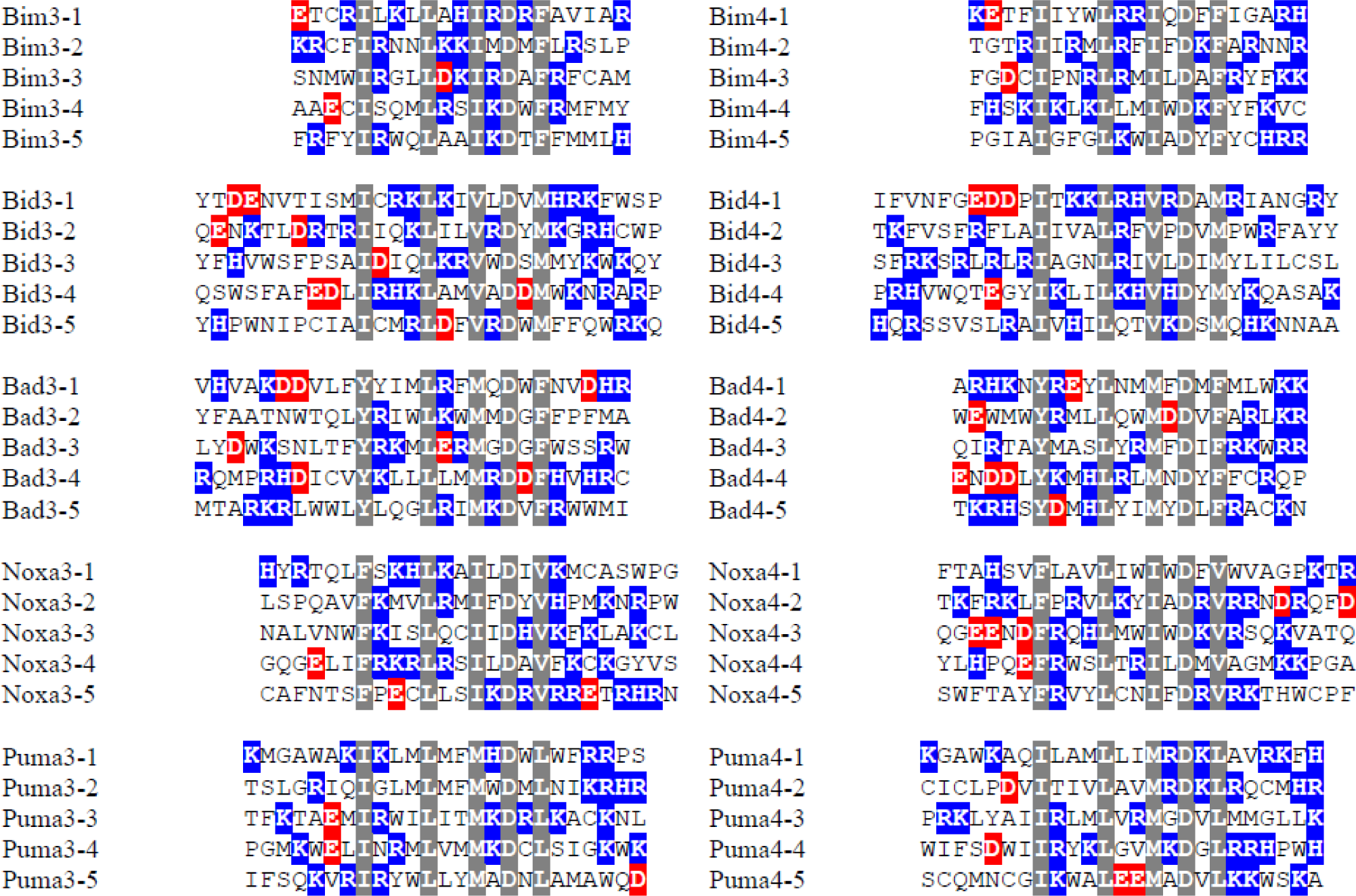
Top BH3-like sequences derived from the wild-type BH3 peptides with highly favorable interaction energies with Bcl-X_L_. Interaction energies of these peptides lie at the negative tail end of the histograms shown in Figure 2 and Figure S1. Those peptides with higher helical propensities and larger hydrophobic moments are selected among the BH3-like sequences that have the most favorable interaction energies with Bcl-X_L_. The four conserved hydrophobic residues and the conserved Asp residue are shown in gray background. The acidic and basic residues are shown in red and blue backgrounds respectively. The sequences shown on the left and right sides correspond to those from Set-I and Set-II respectively.

By looking at the peptide sequences, it is immediately clear that the BH3-like sequences with highly favorable interaction energies with Mcl-1 have more acidic residues (Figure 3). Conversely, the BH3-like sequences tend to have more basic residues when interacting with Bcl-X_L_ and the resultant interaction energies are more favorable than the wild-type BH3 peptides (Figure 4). Helical wheel projections of the wild-type BH3 peptides and the BH3-like sequences derived from the wild-type BH3 peptides are shown in Figures S6 to S10. Vdw component of interaction energies for both Mcl-1 and Bcl-X_L_ complexes for both sets of selected top BH3-like sequences are shown respectively in Figure S10 and Figure S11 respectively. Total interaction energies and the electrostatic component of interaction energies for Mcl-1 and Bcl-X_L_ complexes are shown in Figure 5 and Figure 6 for Set-I. The same data for BH3-like sequences from Set-II are provided in Figures S13 and S14 respectively for Mcl-1 and Bcl-X_L_ complex structures.

**Figure 5:**
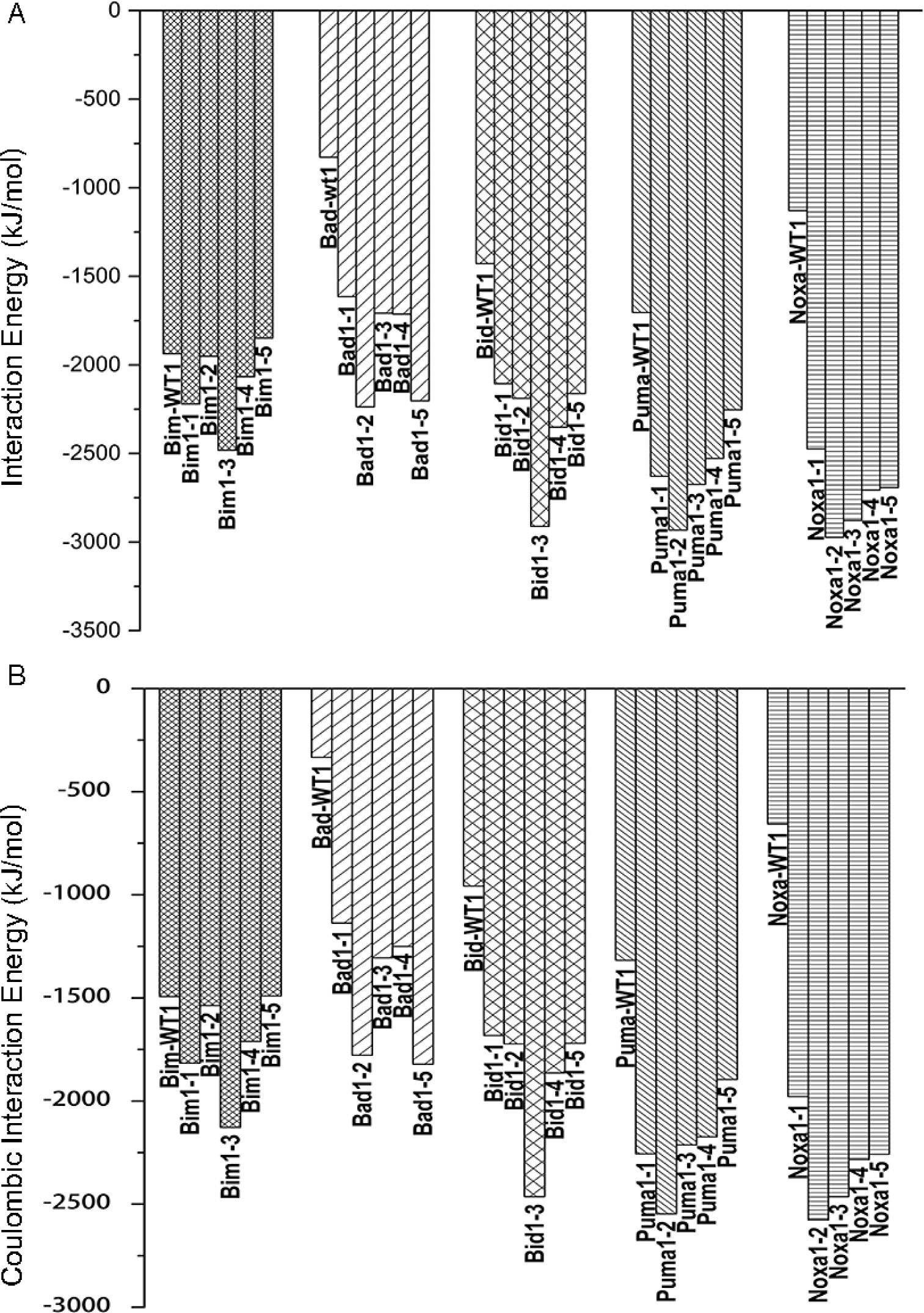
(Top) Interaction energies of wild-type BH3 peptides and top BH3-like peptides from Set-I in complex with Mcl-1. (Bottom) Electrostatic component of interaction energies of wild-type BH3 peptides and top BH3-like peptides from Set-I in complex with Mcl-1

**Figure 6:**
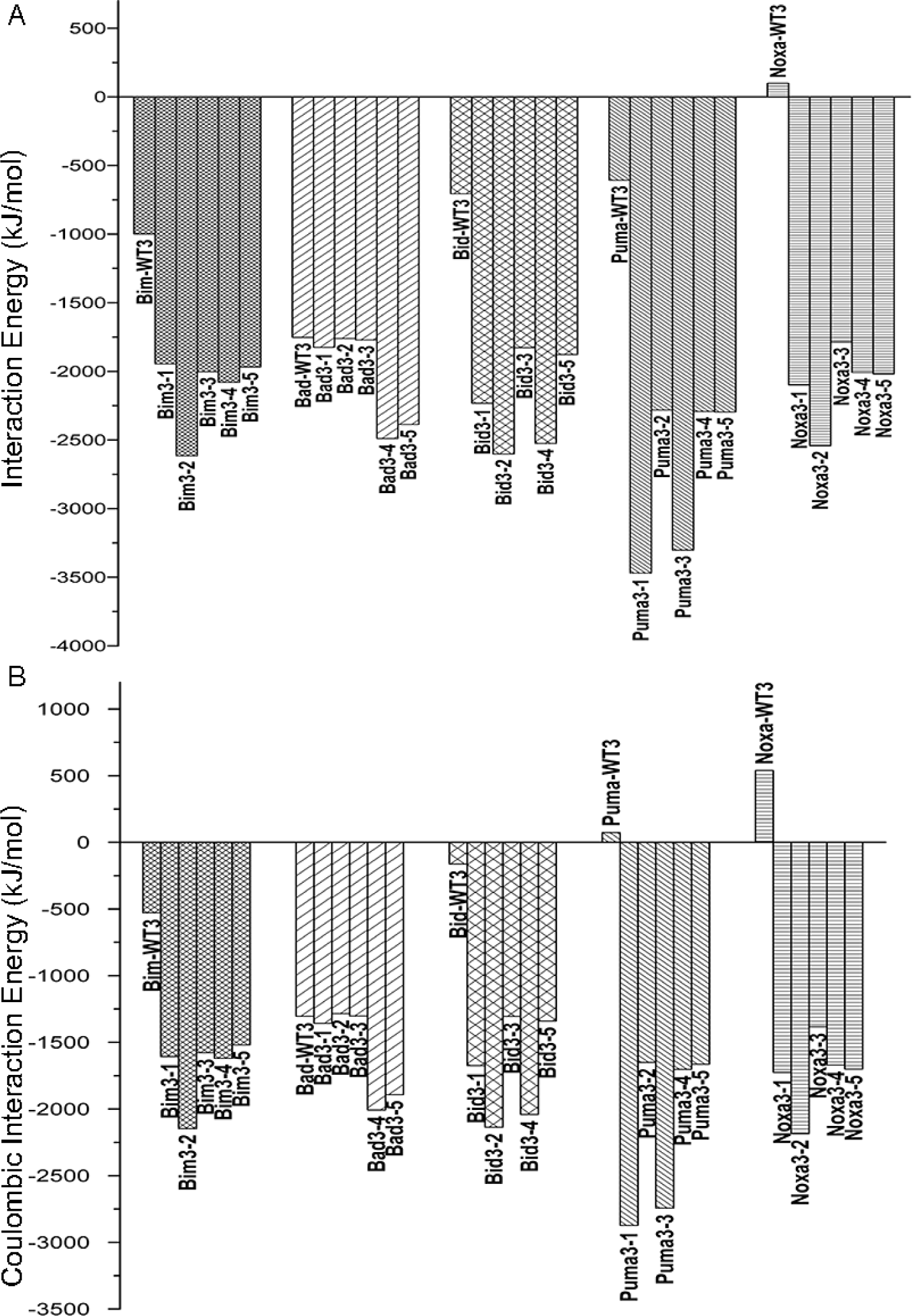
(Top) Interaction energies of wild-type BH3 peptides and top BH3-like peptides from Set-I in complex with Bcl-X_L_. (Bottom) Electrostatic component of interaction energies of wild-type BH3 peptides and top BH3-like peptides from Set-I in complex with Bcl-X_L_.

#### Bim wild-type BH3 and Bim-based BH3-like sequences

Bim wild-type peptides have 2 to 4 basic residues and 3 acidic residues (Figure 1). The selected top Bim-based BH3-like sequences with more favorable interaction energies with Mcl-1 have 3 to 6 acidic residues and 0 to 3 basic residues (Figure 3) and the net charge for the majority of the selected peptides varies from −3 to −6. In the interaction energy histograms, the wild-type Bim BH3 peptides when bound to Mcl-1 have interaction energies close to the negative tail of the distribution (Figure 2 and Figure S1). Interaction energies for the top BH3-peptides with Mcl-1 (Figure 5 and Figure S13) also reveal that they are close to the wild-type Bim BH3 peptide indicating that the sequences of wild-type peptides are already optimized to bind with most favorable interactions with the Mcl-1 protein. This could be attributed to the presence of acidic residues in the wild-type Bim BH3 peptides at the hydrophobic-hydrophilic boundary of the BH3 amphipathic helix that is also the protein-peptide binding interface (Figure S6). The selected Mcl-1-binding Bim-based BH3-like peptides have alpha-helical propensity higher than the Bim wild-type BH3 peptides (Figure S2). The hydrophobic moment of these BH3-like peptides is comparable to the wild-type BH3 peptide (Figure S4). When in complex with Mcl-1, vdw interaction energies of Bim wild-type BH3 peptides (Figure S11) are slightly more favorable than that of Bim-based BH3-like sequences. However, the electrostatic interactions for the Bim-based BH3-like peptides are similar or more favorable to the wild-type Bim BH3 peptides (Figure 5).

The picture with Bcl-X_L_-binding BH3 peptides is, however, quite different. The more favorably interacting Bim-based BH3-like sequences have 4 to 6 basic residues and 0 to 1 acidic residue (Figure 4). The net charge for the selected peptides varied from +2 to +5. Interaction energies of wild-type Bim-BH3 peptides lie close to the average values in the histograms generated from Bim-based BH3 sequences (Figure 2 and Figure S1) when they bind to Bcl-X_L_. This implies that the chance of any random sequence of similar length with the conserved hydrophobic and aspartic residues in equivalent positions that can bind to Bcl-X_L_ with interaction energy of similar magnitude is very high. The top five Bim-derived BH3-like sequences from each library obtained from the negative tail regions interact with Bcl-X_L_ much more favorably than the wild-type Bim peptides (Figure 6 and Figure S14). Helical propensities of these Bim-based BH3-like sequences binding to Bcl-X_L_ are higher than that of the Bim wild-type BH3 sequences (Figure S3). Mean hydrophobic moments of these peptides are either comparable or higher than that found for the wild-type Bim BH3 peptide helices (Figure S5). The presence of several basic residues in the Bim-derived BH3 peptides seems to be responsible for highly favorable interactions with Bcl-X_L_ and it is evident from the interaction energy histograms (Figure 6 and Figure S14). The vdw interaction energies between Bim BH3 wild-type and BH3-like sequences in complex with Bcl-X_L_ are very similar (Figure S12). The major differences are due to the electrostatic interaction energies which are 1000 to 1500 kJ/mol more favorable for the BH3-like sequences than that of the Bim wild-type BH3 peptides (Figure 6 and Figure S14).

#### Bid wild-type BH3 and Bid-based BH3-like sequences

Bid wild-type BH3 peptides have almost equal number of acidic and basic residues (Figure 1). Histograms of interaction energies calculated for the BH3-like sequences derived from Bid wild-type peptides show similar negatively skewed extreme value distributions when binding to Mcl-1 or Bcl-X_L_ (Figure 2 and Figure S1). The wild-type Bim BH3 peptides have interaction energies close to the average values of the distributions in both Mcl-1 and Bcl-X_L_. In Mcl-1 complexes, the top BH3-like sequences with high helical propensities (Figure S2) selected from the negative tail region of the distributions have 4 to 9 acidic residues and 2 to 5 basic residues (Figure 3). The net charge of majority of these peptides is −1 to −5. Helical wheel projections of these top Bim-derived BH3 sequences are shown in Figure S7. While the vdw interaction energies of the Bid-derived BH3 sequences are slightly less favorable than that of wild-type Bid BH3 peptides (Figure S11), the total and electrostatic interaction energies are highly favorable by more than 1000 to 1500 kJ/mol (Figure 5 and Figure S13). While the helical propensities of many of the Bid-based BH3 sequences are comparable to the wild-type Bid BH3 peptides, the hydrophobic moments indicate that wild-type BH3 peptides are likely to form better amphipathic helices (Figure S4).

The top Bid-derived BH3 sequences obtained from the negative tail region of interaction energy histograms bind to Bcl-X_L_ with interaction energies that are > 2000 kJ/mol more favorable than the wild-type Bid BH3 peptides (Figure 2, Figure S1, Figure 6 and Figure S14). While the vdw interaction energies are comparable with Bid wild-type BH3 peptide (Figure S12), electrostatic interactions between the basic residues of Bid-derived BH3 peptides and Bcl-X_L_ are the major contributing factor for the BH3-like sequences obtained from both libraries (Figure 6 and Figure S14). These Bid-based BH3 sequences possess 2 to 4 acidic residues and 4 to 8 basic residues with net charge ranging from +2 to +6 (Figure 4). The helical propensities of BH3-like sequences are higher or comparable to that of the Bid wild-type BH3 peptides (Figure S3). The hydrophobic moments calculated for the BH3-like sequences show that the potential to form an amphipathic helix is predicted to be better for Bid wild-type BH3 peptides (Figure S5 and Figure S7).

#### Bad wild-type BH3 and Bad-based BH3-like sequences

Bad wild-type BH3 peptides do not bind to Mcl-1 ^14,30-31^ and hence we had to model the complex structure of Mcl-1:Bad (see Methods section). As in the case of Bid, Bad wild-type BH3 peptide also possesses almost equal number of acidic and basic residues (Figure 1). Interaction energies calculated for the Mcl-1-bound BH3-like sequences derived from the Bad wild-type peptides exhibit negatively skewed extreme value distributions (Figure 2 and Figure S1). We found similar observation for Bcl-X_L_-bound Bad-based BH3-like sequences also. However in the case of Mcl-1, interaction energies of Bad wild-type peptides are closer to the average interaction energies obtained for all Bad-derived BH3-like peptides. When Bad BH3 wild-type peptides bind to Bcl-X_L_, their interaction energies lie close to the negative tail of the histograms of interaction energies indicating that Bad BH3 wild-type sequences are already optimal to produce highly favorable interaction energies with Bcl-X_L_ (Figure 2 and Figure S1).

The top Mcl-1-binding BH3-like sequences obtained from the negative tails of interaction energy distributions have 3 to 5 acidic residues and 0 to 5 basic residues (Figure 3). The net charge of these peptides varies from −5 to +2. Helical propensities of all the BH3-like sequences are higher than that of wild-type Bad BH3 sequences (Figure S2). Mean hydrophobic moments of many of the Bad-based BH3-like sequences are comparable to that of Bad wild-type BH3 sequences (Figure S4) and the helical wheel projections confirm this (Figure S8). Interaction energies of Bad-based BH3-like sequences with Mcl-1 are more favorable by −500 to −1400 kJ/mol when compared to wild-type Bad BH3 peptides (Figure 5 and Figure 13). With very little difference in the van der Waals component (Figure S11), the most significant difference comes from the electrostatic component (Figure 5 and Figure S13).

The top BH3-like peptides selected from the histogram of interaction energies have 1 to 4 acidic residues and 2 to 8 basic residues when they bind to Bcl-X_L_ (Figure 4) with the net charge ranging mostly from +1 to +5. With strong helical propensities and hydrophobic moments comparable with that of Bad wild-type BH3 peptides (Figure S3 and Figure S5), these Bcl-X_L_-bound Bad-based BH3 sequences exhibit interaction energies more favorable to the Bad wild-type BH3 peptides (Figure 6 and Figure S14). However, when the net charge is +4 or +5 (Bad1-5, Bad2-1 and Bad2-3), they bind to Bcl-X_L_ with interaction energies more than −700 to −1300 kJ/mol significantly stronger than the corresponding wild-type Bad BH3 peptides. As in previous cases, the presence of more positively charged residues facilitate in enhancing the electrostatic interactions of BH3-like sequences with the Bcl-X_L_ protein (Figure 6 and Figure S14) with van der Waals component of interaction energy very similar among all the selected top Bad-derived BH3-like sequences.

#### Noxa BH3 wild-type and Noxa-based BH3-like sequences

In the case of Noxa, experimental studies have shown that its BH3 peptides do not bind to Bcl-X_L_ ^14,31^. Hence, no crystal structure of Bcl-X_L_:Noxa complex is available and as a result, we had to resort to homology modeling to build Bcl-X_L_:Noxa complex structure. As in the previous cases, Noxa wild-type peptide contains nearly equal number of acidic and basic residues (Figure 1). Interaction energies of the BH3-like sequences derived from wild-type Noxa BH3 peptide that bind to Mcl-1 or Bcl-X_L_ are plotted in histograms and the resulting distributions can be described as negatively skewed extreme value distributions (Figure 2 and Figure S1). The wild-type Noxa BH3 peptide has interaction energy close to the average value when it binds to Mcl-1. However, interaction energy between Bcl-X_L_ and Noxa BH3 wild-type peptide is towards the positive side of the histograms indicating that these complexes are energetically not favorable.

The top five Mcl-1-binding Noxa-based BH3-like sequences have 4 to 8 acidic residues and 1 to 4 basic residues with the net charge ranging from −1 to −6 (Figure 3). The helical propensities (Figure S2) and the mean hydrophobic moments (Figure S4 and Figure S9) of these peptides are comparable to wild-type Noxa BH3 peptide. Interaction energies of Noxa-based BH3-like sequences are −1300 to −1800 kJ/mol more favorable than that calculated for the wild-type Noxa BH3 peptide (Figure 5 and Figure S13). In the case of Noxa2-4, it has a net charge of −6 and when it binds to Mcl-1, its interaction energy is more than −2000 kJ/mol favorable than that of Noxa wild-type BH3 peptide. The differences between vdw interaction energies of BH3-like Noxa-derived sequences and the wild-type Noxa BH3 peptide are comparable (Figure S11). Major differences are observed in the case of electrostatic interaction energies. They contribute significantly to the more favorable interactions for the BH3-like peptide sequences (Figure S5 and S13).

The selected top Noxa-based BH3-like sequences from the two libraries when bound to Bcl-X_L_ have 1 to 4 acidic residues and 3 to 9 basic residues with net charge varying from +1 to +6 (Figure 4). Helical propensities (Figure S3) and the mean hydrophobic moments (Figure S5 and Figure S9) of the selected BH3-like sequences are comparable to the wild-type Noxa BH3 peptide. The Noxa wild-type peptide interacts with Bcl-X_L_ with positive interaction energy implying that the wild-type Noxa BH3 peptide residues do not interact favorably with the Bcl-X_L_ protein. However, the Noxa-based BH3-like sequences interact with Bcl-X_L_ favorably with interaction energies ranging from −1700 to −2700 kJ/mol (Figure 6 and Figure S14). Here again we observe that when the net charge is above +4, the interaction energies of the BH3-like peptides with Bcl-X_L_ are −2542 (Noxa3-2) and −2773 (Noxa4-2) kJ/mol indicating that the presence of positively charged residues result in some of the most favorable interactions between the peptide and protein. As anticipated, the electrostatic interactions dominate the total interaction energy between the Noxa-based BH3 peptide and the Bcl-X_L_ protein compared to the vdw component (Figure S12).

#### Puma wild-type BH3 and Puma-based BH3-like sequences

Top BH3-like sequences derived from Puma with highly favorable interaction energies show a very clear bias for acidic residues when interacting with Mcl-1 (Figure 3). These BH3-like peptides have a net negative charge ranging from −3 to −7. Their helical propensities (Figure S2) and mean hydrophobic moments (Figure S4) are mostly comparable with that of wild-type Puma BH3 peptides. Helical wheel projections of top Puma-based BH3-like peptides also present a similar picture (Figure S10). Histograms describing the distribution of interaction energies show that the wild-type Puma BH3 peptides’ interaction energy with Mcl-1 is close to the average of the distribution (Figure 2 and Figure S1) indicating that finding a peptide sequence with similar interaction energy is highly probable. However, interaction energies of top Puma-based BH3 sequences with highly favorable interactions are found close to the negative tail of the distribution. The predominant contribution for the favorable interactions comes from the electrostatic interactions due to the presence of large number of acidic residues (Figure 5 and Figure S13). The peptide Puma2-3 has zero basic residues and seven acidic residues and the electrostatic component of interaction energy between Mcl-1 and this peptide is more than twice favorable than that of wild-type peptide. VDW interactions between the wild-type Puma BH3 peptide and Puma-derived BH3-like peptides are comparable (Figure S11).

The Puma-based BH3 peptides with most favorable interaction energies with Bcl-X_L_ are rich in basic residues (Figure 4). Most of them have net positive charge ranging from +3 to +6. The calculated helical propensities (Figure S3) and the mean hydrophobic moments of most of these peptides (Figure S5) are comparable to that of wild-type Puma BH3 peptides. Interaction energy histograms reveal that interaction energy of the wild-type Puma peptide is close to the average value (Figure 2 and Figure S1). There is a clear distinction between the interaction energies of the wild-type Puma peptided and the Puma-derived BH3 peptides (Figure 6 and Figure S14). While the Puma wild-type BH3 peptides have interaction energies that are little more than −600 kJ/mol, interaction energies of Puma-derived BH3 peptides vary between −2200 to −4100 kJ/mol indicating that Puma-based BH3 peptides interact with Bcl-X_L_ much more favorably than the wild-type Puma peptides. The differences are mainly due to the electrostatic component (Figure 6 and Figure S14). Large number of basic residues in Puma-derived BH3 peptides interacts very strongly with the Bcl-X_L_ protein. There is little difference in the vdw component of interaction energies between the wild-type and Puma-derived BH3 peptides (Figure S12).

### Charged residues from BH3-like sequences distinguish between Mcl-1 and Bcl-X_L_

Results obtained from the BH3-like sequences derived from the BH3 regions of five pro-apoptotic proteins in complex with Mcl-1 or Bcl-X_L_ clearly reveal that BH3-like peptides with high abundance of negatively charged residues interact more favorably with Mcl-1. In the case of Bcl-X_L_, the BH3-like sequences enriched with positively charged residues exhibit highly favorable interaction energies. We have also shown that major contribution of interaction energy between the BH3-like peptide and Mcl-1/Bcl-X_L_ comes from the electrostatic component and vdw component seems to play a minor role. It is interesting to note that a combination of modeling and molecular mechanics approach has clearly highlighted how BH3-like sequences distinguish between Mcl-1 and Bcl-X_L_. We have further extended this study by choosing two representative BH3-like sequences and the corresponding wild-type peptides. We performed molecular dynamics simulations and wet lab experiments to show that these BH3-like peptides do discriminate clearly between Mcl-1 and Bcl-X_L_. Results of the simulation and experimental studies are presented below.

### Molecular Dynamics Simulations of protein-BH3 peptide complexes

To further understand the discriminating nature and strength of interactions of the BH3-like peptides predicted to interact strongly either with Mcl-1 or Bcl-X_L_, we performed molecular dynamics simulations of two of the representative BH3-like peptide sequences derived from the wild-type Noxa (Noxa3-1) and Bad (Bad2-1) in complex with Mcl-1 or Bcl-X_L_. We have chosen Noxa and Bad because the BH3 wild-type peptides of Noxa and Bad do not bind to Bcl-X_L_ and Mcl-1 respectively ^14,30-31^. However in the current study, we have predicted that the Noxa and Bad-derived BH3 peptides, Noxa3-1 and Bad2-1, will bind strongly to Bcl-X_L_ and Mcl-1 respectively. Noxa3-1 and Bad2-1 have respectively several basic and acidic residues (Figure 3 and Figure 4). The wild-type Noxa BH3 peptide binds to Bcl-X_L_ with positive interaction energy and the electrostatic component of interaction energy is not favorable (Figure 6).

Similarly, the wild-type Bad peptide binds to Mcl-1 with interaction energy 700 kJ/mol less favorable than Bad2-1 (Figure 5). Interaction energies between the peptide and the protein were calculated from the minimized protein-peptide complex. It is possible that some of these interactions are weak and less stable and interactions also depend upon the overall helical stability of the interacting BH3 peptides. To investigate these aspects, we have performed several molecular dynamics simulations of the protein-peptide complexes involving Mcl-1/Bcl-X_L_ and Bad2-1/Noxa3-1 BH3 peptides. For comparison, we also performed the simulations with the wild-type Noxa and Bad BH3 peptides in complex with Mcl-1 or Bcl-X_L_. Molecular dynamics protocol that we followed in this study is described in detail in the Supplementary Information. Summary of all the MD simulations performed are provided in Table S1.

We first compared the simulation results of Mcl-1 in complex with the Bad-WT1 and Bad2-1. Three independent simulations of Bad-WT1 in complex with Mcl-1 demonstrated that the Bad BH3 helix significantly lost its helical character at the end of the 500 ns production run in two out of three simulations (Figure S15A-C). Mcl-1 in complex with Bad2-1 however retained most of the helical character in both the simulations (Figure S15D-E). In the case of Noxa3-1, a Noxa-derived BH3-like peptide, a significant loss of helical character was observed in one of the two simulations when this peptide was simulated in complex with Mcl-1 (Figure S16D-E). Our methods predict a strong binding of Noxa3-1 with Bcl-X_L_ and not Mcl-1. Noxa3-1 has six basic residues and one acidic residue. While BH3-like peptides predicted to bind Mcl-1 with high affinity are rich in acidic residues, Noxa3-1 with a net positive charge of +5 is predicted to have poor-binding affinity with Mcl-1. Experimental studies have shown that Noxa wild-type BH3 peptide binds to Mcl-1 strongly ^14,31^. To compare this, we performed three independent simulations of Noxa-WT1 in complex with Mcl-1. In two out of three simulations, the peptide remains helical and strongly bound to the protein (Figure S16A-C). We calculated the electrostatic component of interaction energies between acidic and basic residues of wild-type BH3 or BH3-like peptides with Mcl-1 (Figure 7). In all the ten simulations, the acidic residues have favorable electrostatic interactions with Mcl-1. In Noxa-WT1 and Bad-WT1, basic residues are present. The basic residues of Noxa-WT1 have positive electrostatic interaction energy with Mcl-1. Only in Bad-WT1, basic residues exhibit favorable electrostatic interactions with Mcl-1. The overall electrostatic component of interaction energy between Mcl-1 and Bad-WT1 is favorable and this is achieved in those simulations at the cost of Bad BH3 peptide’s helical character (Figure S15A and S15B). The Bad2-1 peptide exhibited significant favorable electrostatic interactions due to the interactions of its acidic residues with Mcl-1.

**Figure 7:**
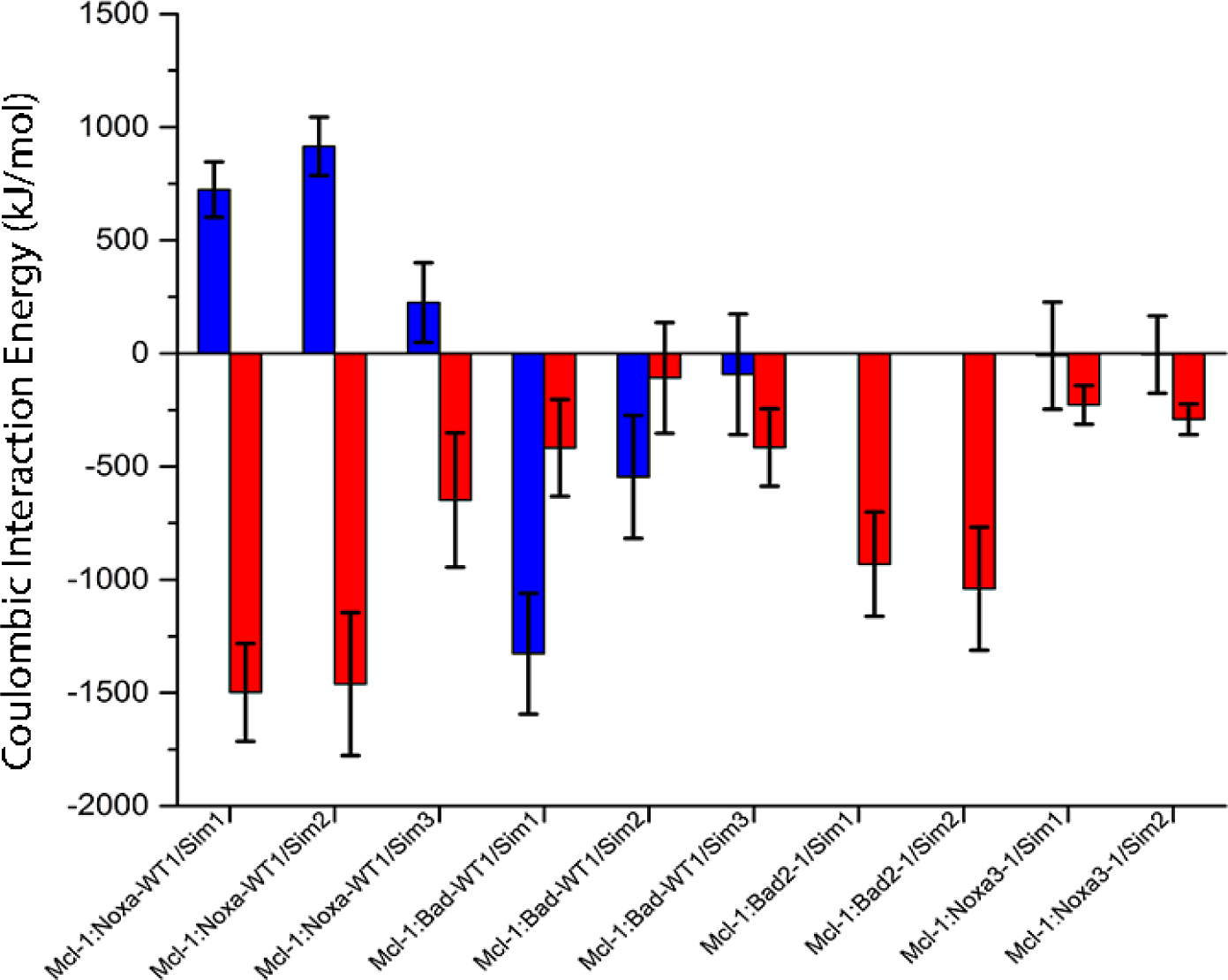
Average electrostatic interaction energies of Mcl-1 in complex with wild-type BH3 peptides Noxa and Bad and BH3-like peptides derived from the wild-type peptides. Electrostatic interactions due to acidic (red) and basic (blue) residues of the BH3 or BH3-like peptides with the Mcl-1 protein are shown separately.

With the exception in one of the simulations of Bad2-1, all BH3 or BH3-like peptides retain helical character to a significant extent when simulated in complex with Bcl-X_L_ (Figure S17 and Figure S18). However, analysis of electrostatic component of interaction energies due to the presence of acidic and basic residues clearly revealed that the basic residues contribute significantly to the favorable nature of interactions with Bcl-X_L_ protein (Figure 8). When acidic residues are present, they result in positive electrostatic interaction energy in all the cases. This is also evident when we compare Bad-WT1 and Bad2-1 and a huge difference in the electrostatic component of interaction energy can be observed between these two peptides. While the electrostatic component of interaction energy for Bad-WT1 is the most favorable that contributes to the overall favorable interaction energy with the Bcl-X_L_ protein, due to the absence of any basic residue, overall electrostatic component of Bad2-1 peptide’s interaction with Bcl-X_L_ is positive. MD simulations of Mcl-1 and Bcl-X_L_ proteins in complex with wild-type Noxa/Bad BH3 peptides or Noxa/Bad-derived BH3-like peptides further reiterate that acidic and basic residues clearly discriminate Mcl-1 and Bcl-X_L_ proteins.

**Figure 8:**
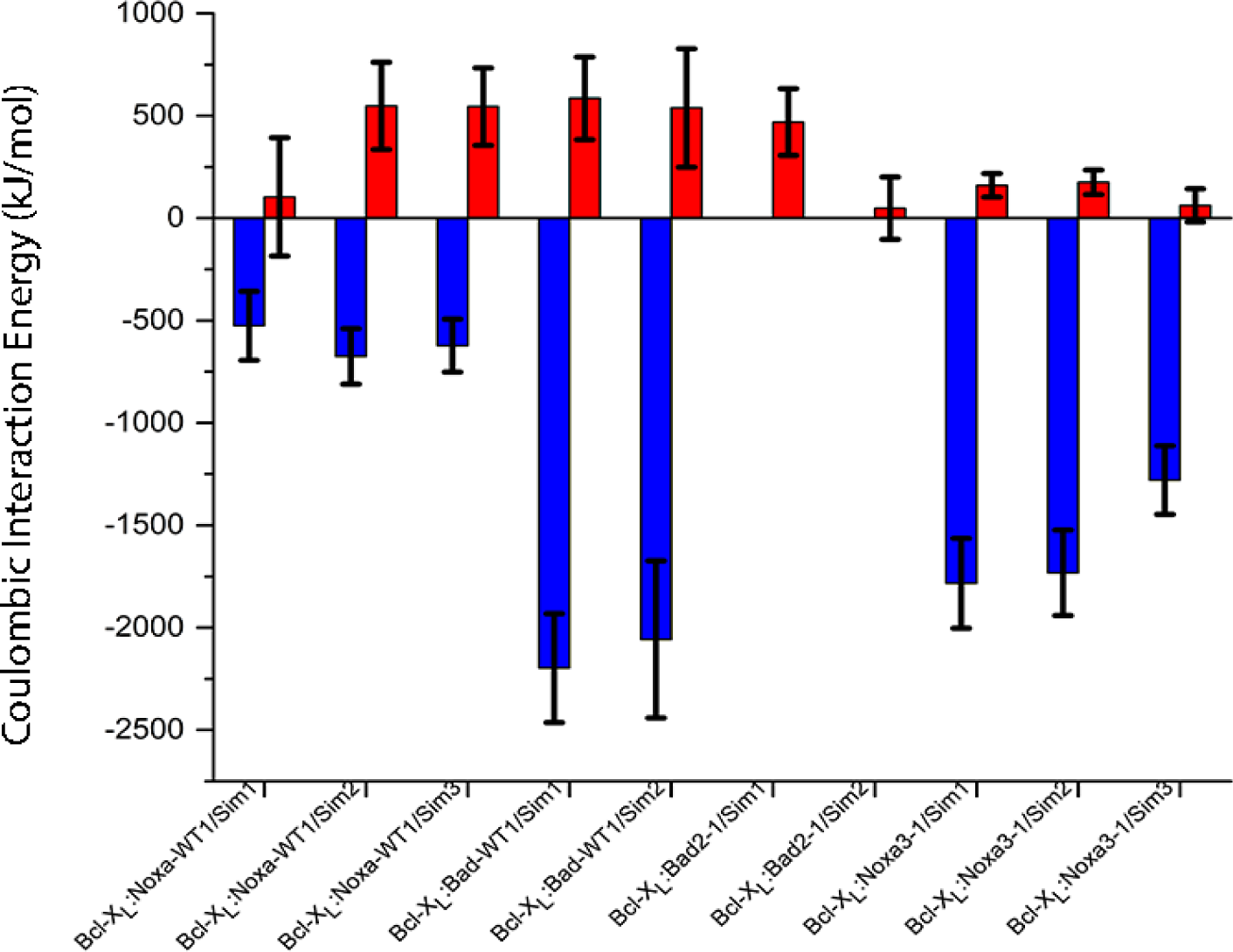
Average electrostatic interaction energies of Bcl-X_L_ in complex with wild-type BH3 peptides Noxa and Bad and BH3-like peptides derived from the wild-type peptides. Electrostatic interactions due to acidic (red) and basic (blue) residues of the BH3 or BH3-like peptides with the Bcl-X_L_ protein are shown separately.

### Synthetic BH3-like peptides hinder the growth of cancer cells

To ascertain the effect of wild-type and BH3-like peptides in vitro, we synthesized four peptides, namely: Noxa-WT1, Bad-WT1, Noxa3-1 and Bad2-1 (Figure 1, 3 and 4). Firstly, we investigated the levels of Mcl-1 and Bcl-X_L_ in prostate cancer PC3 and breast cancer MCF7 cells. We observed higher levels of Mcl-1 in MCF7 as compared to PC3, while elevated expression of Bcl-X_L_ was found in PC3 (Figure 9A). Therefore, we treated the MCF7 cells with two different concentrations (100nM and 500nM) of Noxa-WT1 and Bad2-1 and compared with control (CTL) in which MCF7 cells were not treated with any peptide. We found a marked decrease in the cell viability in the cells treated with both Noxa-WT1 and Bad2-1 peptides. However, treatment with higher concentration of Bad2-1 peptide resulted in ∼60% reduction in the cell viability (Figure 9B). Similarly, PC3 cells treated with Bad-WT1 and Noxa3-1 peptides also exhibit decrease in cell viability, while ∼50% reduction in the cell viability was found with high concentration of Noxa3-1 (Figure 9C). There is no significant difference in cell viability between Noxa3-1 and Bad-WT1 in MCF7 cells (Figure S19A) and this is independent of the concentration used. Similarly, cell viability remained same for both Bad2-1 and Noxa-WT1 in PC3 cells (Figure S19B). We also examined the rate of cell proliferation in these cell lines, and similar to cell viability assay with Bad2-1 and Noxa3-1. Bad2-1 and Noxa3-1 peptide treatment resulted in a significant reduction in the cell proliferation of MCF7 and PC3 respectively (Figure 9D and 9E). Since, cleavage of Poly(ADP-ribose) polymerase (PARP-1) serves as a hallmark of apoptosis ^32^, we next examined whether the reduction in cell viability is due to the activation of apoptotic pathway. We observed increased levels of cleaved PARP in the peptide treated MCF7 and PC3 cells, which suggested that both the Noxa-WT1 and Bad2-1 are leading to induction of apoptotic pathway in MCF7 cells (Figure 9F). A similar observation was made for Bad-WT1 and Noxa3-1 in PC3 cells.

**Figure 9:**
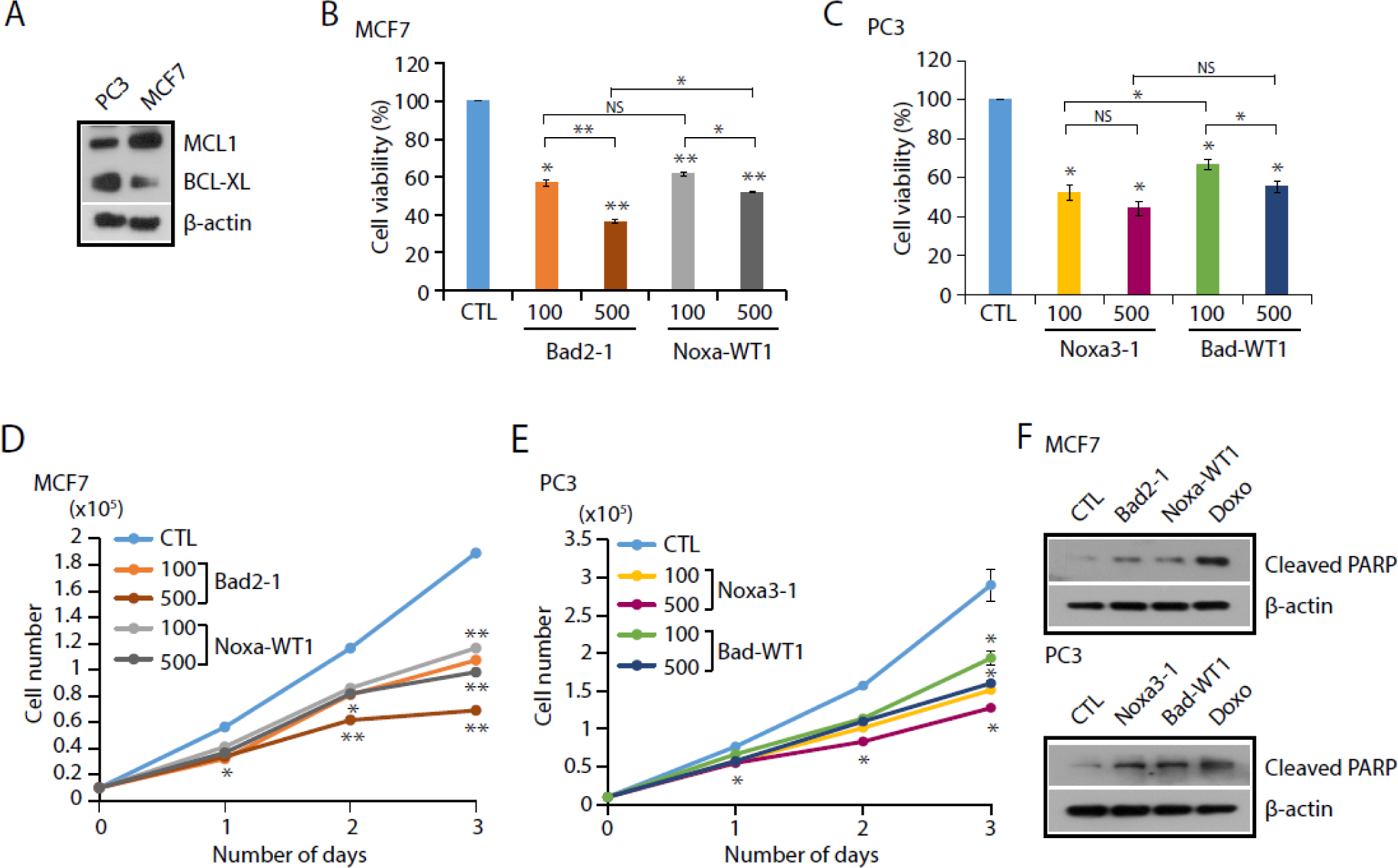
Synthetic BH3-like peptides against MCL1 and BCL-X_L_ induce apoptosis in cancer cells. (**A**) Immunoblot for MCL1 and BCL-X_L_ levels in MCF7 and PC3 cells; β-actin was used as loading control. (**B**) Percent cell viability using MCF7 cells treated with different concentrations (100nM and 500nM) of Bad2-1 and Noxa-WT1 peptides along with control (CTL). (**C**) Percent cell viability using PC3 cells treated with different concentrations (100nM and 500nM) of Noxa3-1 and Bad-WT1 peptides along with CTL. (**D**) Cell proliferation assay using same cells as in (B). (E) Cell proliferation assay using same cells as in (C). (**F**) Immunoblot for cleaved PARP levels in MCF7 cells treated with Bad2-1 and Noxa-WT1 peptides (500nM) along with CTL (Top). Immunoblot for cleaved PARP levels in PC3 cells treated with Noxa3-1 and Bad-WT1 peptides (500nM) along with CTL (Bottom).

## Discussion

Anti-apoptotic Bcl-2 proteins have been the targets of cancer therapy for many years and efforts to develop efficient inhibitors have resulted in many promising compounds ^1,9,15-20^. One of the strategies is to use the BH3 peptides of pro-apoptotic Bcl-2 proteins to antagonize the anti-apoptotic Bcl-2 members and thus cellular death pathway can be activated in cancer cells ^24-25,27^. Although several pro-apoptotic BH3 peptides have affinity for anti-apoptotic proteins at nM level, the BH3 peptide inhibitors have cross reactivity with many anti-apoptotic Bcl-2 partners ^14,13^. Efforts to synthesize BH3 mimetic peptides that have stable helical structure and enhanced affinity for specific anti-apoptotic Bcl-2 proteins have yielded mixed results ^23-26,28,33^. These studies focused mainly on the hydrophobic face of the amphipathic BH3 helix. However, many studies have shown that in addition to the interaction of hydrophobic face of pro-apoptotic BH3 amphipathic helix with the exposed hydrophobic patch of anti-apoptotic Bcl-2 protein, hydrophilic interactions between specific residues play a major role in giving specificity to the pro-apoptotic Bcl-2 proteins ^34-36^. Techniques such as FACS and deep sequencing were used to obtain high-throughput data on protein-peptide binding ^28^. As part of the strategy to use structure-based design, high-affinity peptide-binders were designed based on “recurring tertiary structural motifs” from known protein structures available in PDB and this was applied to Bcl-X_L_, Mcl-1 and Bfl-1 proteins ^29^. Staples were introduced that enhanced the stability of α-helical structure of peptides that inhibited anti-apoptotic proteins with increased affinity ^24,37-38^. In this paper, we have developed a novel computational method to generate BH3-like sequences that can bind Mcl-1 or Bcl-X_L_ with high affinity and specificity.

### Computational design of BH3-like peptides

The method developed in this study to design BH3-like sequences is inspired by the approach used to determine the statistical significance of pairwise sequence alignment of two protein/DNA sequences ^39-40^. In this process, the distribution of alignment scores obtained from shuffled sequences result in a positively skewed Gumbel extreme value distribution. If the two sequences are related, then its score lies in the extreme positive tail of the distribution and the probability of obtaining such a high score purely by chance from a random sequence is extremely low. Using a similar approach, we randomized the sequence of the original BH3 wild-type peptides and only the four conserved hydrophobic residues and the conserved Asp residue (Figure 1) were retained in their original positions. All other residues were substituted randomly to generate a large number of BH3-like sequences. Notably, in earlier studies the conserved hydrophobic residues and the residues at the interface of peptide-protein binding region were substituted to generate the peptide libraries ^23,29^. In our approach, we retained the conserved hydrophobic and Asp residues and randomly substituted residues at all other positions without introducing any bias.

Two sets of 1000 randomized sequences thus obtained for each of the five BH3 wild-type peptides were used to model the protein-peptide complex and both Mcl-1 and Bcl-X_L_ were considered. In the 20000 homology models of BH3-like sequences in complex with Mcl-1 or Bcl-X_L_, the protein structure and the backbone of the peptide helical segment from the template structures were retained. Only the side-chains of the BH3-like peptides were modeled. We hypothesized that the interaction energies of BH3-like peptides in complex with Mcl-1 and Bcl-X_L_ should give a clue about the strength of binding. The stronger the interaction energy between the BH3-like peptide and the Mcl-1/Bcl-X_L_ protein, the stronger will be the binding affinity of the peptide. We determined the interaction energies between the peptide and the protein as described in the Methods section. Distributions obtained for the interaction energies for different sets of BH3-like peptides resulted in negatively-skewed extreme value distributions (Figure 2 and Figure S1). Interaction energies of wild-type BH3 peptides which bind strongly to the protein and are known to have high binding affinity fall in the extreme negative tail end of their respective distributions (for example, see Mcl-1:Bim and Bcl-X_L_:Bad complexes in Figure 1). Interaction energy of Noxa wild-type peptide which is known to have poor binding affinity for Bcl-X_L_ lies towards the positive side of the distribution profile (Figure 1I).

Experimental and computational studies showed a direct relationship between stable helical nature of BH3 peptides and binding affinities ^41-43^. Hence, we selected the peptides from the extreme negative tail of the distributions which are predicted to have high helical propensity and has higher potential to form amphipathic helices. In majority of the peptides, the interaction energies were found to occur beyond mean - 2 standard deviation. In other words, the probability of finding a randomly generated peptide sequence to have such highly favorable interaction energy will be very low. First, we analyzed the chemical nature of these selected peptides. To our surprise, we found that the BH3-like peptides that bind to Mcl-1 strongly are rich in acidic residues (Figure 3). When the same set of BH3-like sequences was used against Bcl-X_L_, the top selected BH3-like peptides from the extreme negative tail that have strong interactions with Bcl-X_L_ are enriched with basic residues (Figure 4). It is also important to note that the acidic/basic residues need not be at the binding site or at the interface of binding region (Figures S6 to S10). The enrichment of the basic or acidic residues seems to be an important criterion that will determine whether a given BH3-like peptide is likely to bind Mcl-1 or Bcl-X_L_. Since we have not changed the conserved hydrophobic residues at the four positions and the conserved Asp residue is also retained, the BH3-like peptide that can form potential amphipathic helix has already the required features to bind the anti-apoptotic Bcl-2 protein. Electrostatic interactions are long range in nature and it appears that the field generated by the acidic or basic residues of the BH3-like peptides participates in long range interactions with the oppositely charged residues in the protein to which they bind. Thus even if the charged residues are not in the peptide-protein binding interface, the long range electrostatic interactions due to these charged residues have significant effect on binding. This observation was confirmed when two independent sets of randomized BH3-like sequences derived from each of five wild-type pro-apoptotic BH3 peptide was tested with Mcl-1 or Bcl-X_L_. Although the previous studies have identified polar interactions between specific residues responsible for high binding affinity or specificity ^34-36^, our studies show that the long-range interactions as a result of overall acidic or basic nature of the peptide can determine the specificity and affinity of BH3-like peptides to bind either Mcl-1 or Bcl-X_L_.

### Experimental validation of the specificity of predicted BH3-like peptides

To validate the computational prediction, we selected two representative BH3-like peptides (Bad2-1 and Noxa3-1) and their corresponding wild-type BH3 peptides (Noxa-WT1 and Bad-WT1). It has been shown in previous studies that Bad-WT1 does not bind to Mcl-1 ^14,31^. However, Bad2-1 drastically decreased the cell viability in concentration-dependent manner in MCF7 cells where Mcl-1 has higher level of expression (Figure 9B). This effect is even more pronounced when compared to Noxa-WT1 indicating that Bad2-1 binds to Mcl-1 with higher affinity than that of Noxa-WT1. The predicted helical propensity of Bad2-1 is higher than that of Bad-WT1 (Figure S2B). The hydrophobic moment of Bad2-1 is comparable to that of Bad-WT1 (Figure S4B). Bad2-1 has no basic residue and has a net charge of −4 while the Bad-WT1 has a net charge of +1 with four basic residues. The absence of basic residues and the increase in the number of acidic residues enabled Bad2-1 to bind Mcl-1 protein strongly even though Bad2-1 has the same conserved hydrophobic and Asp residues at the same positions as that of Bad-WT1. This indicates that the net negative charge of the peptide and by extrapolation the long range attractive electrostatic interactions between Bad2-1 and Mcl-1 result in a tighter binding of Bad2-1 with Mcl-1. Two independent MD simulations of Bad2-1 peptide in complex with Mcl-1 confirm that the average electrostatic interaction energies due to acidic residues is much more favorable compared to the Bad wild-type peptide (Figure 7). The effect of Noxa3-1 on cell viability in MCF7 cells is same as that of Bad-WT1 indicating that Noxa3-1 does not have much affinity for Mcl-1 (Figure S19A).

With the elevated expression of Bcl-X_L_ in PC3 cell lines (Figure 9A), the computationally designed Noxa3-1 results in much reduced cell viability in PC3 cells in concentration-dependent manner and the effect is more pronounced than that of Bad-WT1 (Figure 9C). Noxa3-1 has higher helical propensity than that of Noxa-WT1 (Figure S3A). The hydrophobic moments of both peptides are similar (Figure S5A). While Noxa3-1 has six basic residues and 1 acidic residue (Figure 4), Noxa-WT1 has four acidic residues and three basic residues (Figure 1). Thus the net charge in Noxa3-1 and Noxa-WT1 is +5 and −1 respectively. Although Noxa3-1 has the same conserved hydrophobic residues, unlike Noxa-WT1, Noxa3-1 binds to Bcl-X_L_ and it can be attributed to the presence of many of its basic residues. Three independent simulations carried out on the Bcl-X_L_:Noxa3-1 complex structures indicate that the average electrostatic interaction energies due to the presence of basic residues are highly favorable (Figure 8). The logical conclusion from this observation is that the long-range electrostatic interactions between Noxa3-1 and Bcl-X_L_ play a significant role in binding to the protein. In the Bcl-XL-enriched PC3 cells, cell viability found for Bad2-1 is not significantly different from Noxa-WT1 indicating that Bad2-1 does not bind to Bcl-X_L_ (Figure S19B).

Thus cell viability and cell proliferation studies along with the results of from MD simulations corroborated that acidic and basic residues in BH3-like peptides have more favorable interactions with Mcl-1 and Bcl-X_L_ respectively. The current study has presented a computational method to design BH3-like sequences that can bind specifically to Mcl-1 or Bcl-X_L_. Binding affinities of BH3-like peptides predicted by this method to the specific anti-apoptotic protein can be further improved by introducing staples in suitable positions. Such peptides specific to Mcl-1 or Bcl-XL can be a powerful tool in BH3-profiling assays in which the specific anti-apoptotic Bcl-2 protein that is overexpressed to detect resistance to apoptosis due to Mcl-1 or Bcl-XL. The method proposed here can be easily extended to other anti-apoptotic proteins and viral Bcl-2 homologs.

## Conclusions

In this paper, we have developed a novel computational method to design BH3-like sequences that can bind specifically to Mcl-1 or Bcl-X_L_ with high affinity. BH3-like sequences are generated from wild-type pro-apoptotic BH3 peptides by keeping the four conserved hydrophobic and aspartic residues and substituting all other positions randomly. The peptides generated using this method are then used to model complex structures with Mcl-1 or Bcl-X_L_. Interaction energies calculated between the peptide and the protein give rise to a negatively skewed extreme value distribution. The peptides selected from the extreme negative tail of this distribution have highly favorable interaction energy with the protein. These peptides are rich in acidic residues when they bind to Mcl-1. However, they show distinct preference for basic residues when they are in complex with Bcl-X_L_. The long range interactions due to the charged residues play an important role in highly favorable interaction energies. Cell viability and cell proliferation experiments performed on representative BH3-like sequences clearly demonstrate that the Noxa-derived and Bad-derived BH3-like sequences bind to Bcl-X_L_ and Mcl-1 respectively while it has been shown that the wild-type Noxa and Bad peptides do not bind to Bcl-X_L_ and Mcl-1. The peptides thus obtained can be further improved by introducing staple linkers in suitable positions and/or capping residues to promote helical stability and this will serve as starting point for developing peptide-based BH3-mimetic therapeutic molecules as anti-cancer drugs. This method can easily be extended to other anti-apoptotic Bcl-2 proteins and viral Bcl-2 homologs.

## Materials and Methods

### Generation of BH3-like sequences

We generated BH3-like sequences using the following protocol. For this purpose, we considered BH3 regions of five pro-apoptotic proteins that included Puma, Bim, Bid, Bad and Noxa. Residues corresponding to the four highly conserved hydrophobic positions and the conserved Asp were retained as it is (Figure 1). All the remaining positions were substituted randomly using Sequence Manipulation Suite web server (http://www.bioinformatics.org/sms2/random_protein_regions.html) developed by Paul Stothard^44^. For each BH3-only peptide, we generated two sets (Set-I and Set-II) of 1000 BH3-like sequences.

### Modeling the protein-peptide complex

We used experimentally determined structures of Bcl-X_L_ and Mcl-1 in complex with the BH3 peptides to model the BH3-like sequences bound to Bcl-X_L_ or Mcl-1. The Bcl-X_L_ complex structures used as templates are Bcl-X_L_:Puma (PDB ID: 2M04) ^45^, Bcl-X_L_:Bid (PDB ID: 4QVE; resolution: 2.05 Å) ^46^, Bcl-X_L_:Bad (PDB ID: 2BZW; resolution: 2.3 Å) ^47^ and Bcl-X_L_:Bim (PDB ID: 4QVF; resolution: 1.531 Å) ^46^. For Mcl-1, the complex structures are Mcl-1:Puma (PDB ID: 2ROC) ^48^, Mcl-1:Noxa (PDB ID: 2ROD) ^48^, Mcl-1:Bid (PDB ID: 2KBW) ^49^ and Mcl-1:Bim (PDB ID: 3KJ0; resolution: 1.7 Å) ^30^. The Mcl-1 complex structures with Puma, Noxa and Bid BH3 peptides were determined using solution NMR. Among the many conformers submitted to PDB for the NMR structures, the first model was considered for each of these complex structures. The structures of Mcl-1:Bad and Bcl-X_L_:Noxa have not been determined experimentally. Hence, the native Mcl-1:Bad and Bcl-X_L_:Noxa complex structures were modeled using the experimentally determined Mcl-1:Bim (PDB ID: 3KJ0; resolution: 1.7 Å) and Bcl-X_L_:Bim (PDB ID: 1PQ1; resolution: 1.65 Å) structures respectively as templates. We used MODELLER 9.14 ^50-52^ to model Mcl-1:Bad, Bcl-X_L_:Noxa and all the protein-BH3-like peptide complexes. If there are missing residues or missing loop regions in Mcl-1 or Bcl-X_L_ proteins, they were added using other known Mcl-1 or Bcl-X_L_ apo-or complex structures. The models generated for the BH3-like peptides in complex with Mcl-1 or Bcl-X_L_ will differ only for the BH3-only peptides as the protein is the same for both the template and target. The critical step in homology modeling using MODELLER is the target-template sequence alignment. Since the proteins are the same, the target-template alignment boils down to correctly aligning the BH3 peptide of the wild-type from the template and the BH3-like peptide sequences. Since we have not changed the residues at the four conserved hydrophobic positions and the conserved Asp, target-template alignment of the sequences was straightforward. However, we had to deal with couple of issues that are described below. For a given BH3-like sequence, we considered two different template structures for modeling Mcl-1 and Bcl-X_L_ and the peptide length in these complex structures differed by few residues. Hence, the two sets of BH3-like sequence libraries we generated differed in length. If the sequence length of the BH3 region in a template structure considered for Mcl-1 (or Bcl-X_L_) complex is longer, then the BH3-like sequences that were generated based on the experimentally determined peptide length are also longer. When this library was used to model the complex structure of Bcl-X_L_ (or Mcl-1) and if the corresponding peptide length in the template structure is shorter, then we only considered the sequence that fitted the peptide length in the complex structures and truncated N- and/or C-terminal regions. In the second situation, if the length of BH3-like sequences generated using Mcl-1 (or Bcl-X_L_) template sequence is shorter than the peptide length in the template structure of Bcl-X_L_ (or Mcl-1), then we truncated the peptide length in the template structure in the N- and/or C-terminal region. The wild-type pro-apoptotic BH3 peptides that were used to generate BH3-like sequences and the PDB IDs that were used as templates to model the Mcl-1 and Bcl-X_L_ complex structures are provided in Figure 1.

In MODELLER, the spatial restraints derived from the template structure are represented in the form of an objective function and this objective function is optimized for the target structure. CHARMM22 force filed ^53^ was used for this purpose. For each BH3-like sequence, we used the same modeling protocol to obtain the complex structures with Mcl-1 and Bcl-X_L_. For each complex, the top five best models were considered further for energy minimization.

#### Energy minimization of modeled structures

The top five modeled complex structures for each BH3-like sequence were solvated in a box of water molecules with the protein-peptide complex at the center of the box. SPC/E water ^54^ model was used for this purpose. The entire system was made neutral by adding required number of Na^+^ and Cl^-^ ions. The minimum distance between the protein-peptide complex and the wall of the box was kept at least 10 Å. All-atom force-filed OPLS/AA ^55^ was used to minimize the energy of the system. The same force-field was also used to estimate the interaction energy between the protein and the BH3-like peptide. We used twin-range cut-off to calculate the non-bonded interactions. A cut-off of 10 Å was used to calculate the short-range interactions and the long-range interactions were calculated for the non-bonded interaction pairs that were within 10 to 25 Å. Energy minimization was carried out using GROMACS 4.5.5 ^56^. Among the top five best models, the structure with the most favorable potential energy was chosen to calculate the interaction energy between Mcl-1/Bcl-X_L_ protein and the BH3-like peptide using equation (1).

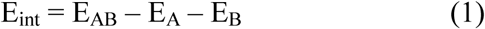

Where E_int_ is the interaction energy between the protein and the BH3-like peptide, E_AB_ is the energy of the protein and BH3-like peptide complex, E_A_ is the energy of the protein and E_B_ is the energy of the BH3-like peptide. Interaction energy is composed of short-range and long-range electrostatic and van der Waals interactions. Two sets of one thousand BH3-like peptide sequences were generated by randomizing each of the five BH3 sequences of Puma, Bim, Bid, Bad and Noxa as explained above. BH3-like sequences derived from the five wild-type BH3 peptides were used to model Mcl-1 and Bcl-X_L_ complex structures. Interaction energies between BH3-like peptides and Mcl-1 or Bcl-X_L_ proteins were calculated for 20,000 modeled complex structures. For comparison purpose, interaction energies of the five wild-type BH3 peptides with Mcl-1 or Bcl-X_L_ were also calculated using the same protocol.

#### Analysis of distribution of interaction energies

For each wild-type BH3 peptide, two sets (Set-I & Set-II) of 1000 randomized BH3-like sequences were generated and the complex structures with Mcl-1 (or Bcl-X_L_) were modeled. Interaction energies between Mcl-1 (or Bcl-X_L_) and the BH3-like peptides were calculated from the modeled complex structures as described above. Histograms describing the distribution of interaction energies were plotted and one such histogram is shown for Bim-based BH3 sequences in complex with Mcl-1 in Figure 2A. It is clear that this is not a normal distribution and is negatively skewed. This distribution can be described as Gumbel extreme minimum value distribution and it is fitted using Matlab R2013 with Levenberg Marquardat algorithm ^57^. From this distribution, we considered only those BH3-like peptides whose interaction energies fall below two to three standard deviations from the mean value. With highly favorable interaction energies, these BH3-like peptides are likely to exhibit higher binding affinity and specificity for that particular anti-apoptotic protein.

#### Helical character and amphipathic nature of BH3-like sequences

BH3 domains of Bcl-2 proteins form amphipathic helices and their helical stability is directly related to their affinities to bind the anti-apoptotic Bcl-2 proteins ^41-43^. The hydrophobic side of the BH3 helix binds to the hydrophobic groove of the binding partners. Hence, we wanted to find out the helix-forming tendency of BH3-like peptides with most favorable interaction energies. We also wanted to determine if they can form helix, whether these helices formed by the BH3-like sequences are likely to be amphipathic in nature. Helical nature of all BH3-like sequences was determined using the AGADIR web server ^58-60^. Helicity was determined at 278 K, pH 7 in 0.1 M ionic strength. Mean helical hydrophobic moment (μH) is a measure of amphiphilicity. Larger the value of μH, higher will be the amphiphilicity. We used HELIQUEST web server ^61^ to determine μH for each BH3-like sequence.

### Cell growth study using BH3-like peptides

#### Cell culture

The prostate cancer cell line (PC3) and breast cancer cell line (MCF7) were procured from American Type Cell Culture (ATCC, Manassas, VA, USA). Both the cell lines were cultured in a CO_2_ incubator (Thermo-Fisher) supplied with 5% CO_2_ at 37°C temperature as per the ATCC guidelines. Short tandem repeats (STR) profiling for cell line authentication was done at the DNA Forensics Laboratory, New Delhi. Mycoplasma contamination of all the cell lines was checked routinely using PlasmoTest mycoplasma detection kit (InvivoGen).

#### BH3-like peptides sequence and synthesis

We synthesized the small peptides (∼45 amino acids) with the wild-type (WT) and BH3-like sequences for NOXA and BAD at GL Biochem, Shanghai Ltd., China. These four peptides contained a N-terminal cell penetrating peptide (TAT) sequence (GRKKRRQRRRPQ) and C-terminal FLAG tag (DYKDDDDK).

Noxa-WT1: GRKKRRQRRRPQ-AELPPEFAAQLRKIGDKVYCTWSAPD-DYKDDDDK

Bad-WT1: GRKKRRQRRRPQ-WAAQRYGRELRRMSDEFEGSFKG-DYKDDDDK

Noxa3-1: GRKKRRQRRRPQ-HYRTQLFSKHLKAILDIVKMCASWPG-DYKDDDDK

Bad2-1: GRKKRRQRRRPQ-DDVMGYEDYLQMMLDFFFAFCN-DYKDDDDK

#### Western blotting assay

Cultured cells were scraped using radioimmunoprecipitation assay (RIPA) lysis buffer, supplemented with complete Protease Inhibitor Cocktail (VWR, M-250) and PhosSTOP (Roche, 4906845001). Protein estimation was done using Bicinchoninic Acid (BCA) Protein Assay kit (GBiosciences, 786-570) and the samples were prepared in 4X SDS-PAGE loading buffer. Protein samples were resolved on the SDS-PAGE and transferred onto a polyvinylidene difluoride (PVDF) membrane (GE Healthcare). The membrane was blocked with 5% non-fat dry milk in tris-buffered saline, 0.1% Tween 20 (TBS-T) for 1 hour at room temperature, and then incubated overnight at 4°C with the following primary antibodies: 1:1000 diluted Mcl-1 (Abcam, ab32087), 1:1000 diluted Bcl-X_L_ (CST, #2764), 1:1000 diluted Cleaved PARP (Asp214) and 1:5000 diluted β-actin (Abcam, ab6276). Subsequently, blots were washed in 1X TBS-T buffer and incubated with respective horseradish peroxidase (HRP) conjugated secondary Anti-Mouse IgG (H+L) (Jackson ImmunoResearch, 715-035-150) or Anti-Rabbit IgG (H+L) (Jackson ImmunoResearch, 711-035-152) antibody for 2 hours at room temperature. For protein detection by Western blot analysis, the signals were visualized using SuperSignal West Chemiluminescent Substrate (Thermo Scientific) following the manufacturer’s protocol. For all immunoblot experiments, β-actin was used as a loading control.

#### Cell viability assay

To determine the effect of the synthesized peptides on viability of prostate cancer and breast cancer cell lines, 3000 cells were plated in each well of the 96-well plate (Nunc, 167008). After 24 hours, synthetic peptides were added to the media at two different concentrations: 100nM and 500nM to the cells, along with no peptide treatment as the control (CTL). After 72 hours of peptide treatment, cell viability was determined using Cell Proliferation Reagent WST-1 (Roche, 5015944001) following the manufacturer’s instructions. Briefly, WST-1 was added to the cultured cells in each well at 1:10 final dilution and incubated for 4 hours at 37°C. The absorbance was measured at 450 nm using a multi-well spectrophotometer (Thermo Scientific Multiskan EX, 51118170); the measured absorbance can be directly correlated to the percent cell viability as per the following equation:

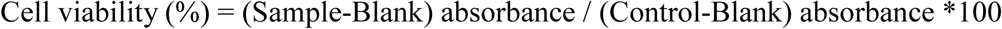

#### Cell proliferation assay

To observe the rate of cell proliferation upon peptide treatment, 10000 cells were plated in each well of a 12-well plate (Corning, 3512). Cell proliferation for the peptide treated cells and no peptide treatment as control (CTL) were measured for three consecutive days using a Beckman Coulter Z2 Particle Count and Size Analyzer (Beckman Coulter, Brea, CA, USA). The number of cells are calculated and plotted for each day.

#### Statistical significance

Biologically independent samples were used (n=3) in each experiment. Error bars represent standard error of the mean (SEM) and data is plotted as mean ± SEM. Statistical significance was determined using two-tailed unpaired Student’s t-test, ∗P≤ 0.05 and ∗∗P≤ 0.001.

### Molecular Dynamics Simulations

We selected two BH3-like peptides Noxa3-1 and Bad2-1 derived respectively from the BH3 regions of Noxa and Bad wild-type proteins. We considered the structures of Mcl-1 and Bcl-X_L_ in complex with Noxa3-1 and Bad2-1 and performed molecular dynamics (MD) simulations. For comparison purpose, we also carried out MD simulations of the same proteins in complex with Noxa-WT1 and Bad-WT1. In each case, the protein-peptide complex structure was placed at the center of a cubic box such that the minimum distance between the complex and the wall was at least 13 Å. The complex was then solvated using SPC water model ^54^ and sufficient amount of Na+ or Cl-ions were added to make the system neutral. OPLS all atom force-field ^55^ was used and LINCS algorithm ^62^ was used to constraint bonds involving hydrogen atoms. After the energy minimization, the system was equilibrated by performing MD simulation in NVT ensemble for 1 ns. This was followed by another round of equilibration in NPT ensemble for about 3.8 ns. Temperature was maintained at 300 K by coupling protein and solvent molecules separately to modified Berendsen thermostat ^63^ with coupling constant of 0.1 ps. Pressure was maintained at 1 atm by coupling Parrinello-Rehman barostat to system at 0.2 ps coupling constant ^64^. During NVT equilibration, positional restraints (force constant 2000 kJ/mol/nm^2^) were applied on the heavy atoms of complex for entire 1 ns simulation. During the initial 1.8 ns of NPT simulation, excluding the backbone heavy atoms of peptide, positional restraints on the heavy atoms of rest of the complex were gradually removed (force constant 2000 kJ/mol/nm2 to 0 kJ/mol/nm2). In the rest of NPT equilibration (2 ns), positional restraints of the backbone heavy atoms of peptide were gradually removed (force constant from 2000 kJ/mol/nm2 to 0 kJ/mol/nm2). This is to ensure that the BH3 peptide helix is properly equilibrated and not destabilized due to any other factors. Production run for each system was carried out for a period of 500 ns and MD-simulated structures were saved for every 10 ps for further analysis. For equilibration and production runs van der Waal interactions were treated using a cut-off of 10 Å and electrostatic interactions were treated using Particle Mesh Ewald method (PME) ^65^.

Interaction energy between the protein and the BH3 peptide was calculated for each MD-simulated structure. A twin-range cut-off of 10 and 25 Å was used by considering peptide and protein as two separate groups.

## Acknowledgements

We thank the High Performance Computing Facility at IIT-Kanpur for making the computational resources available. CNR gratefully acknowledges the BINC Fellowship from the Department of Biotechnology, Government of India. N.M. acknowledges fellowship support from the University Grants Commission (UGC), Government of India. RS is Pradeep Sindhu Chair Professor. BA is thankful for the research funding from the Wellcome Trust/DBT India Alliance (IA/I(S)/12/2/500635), Department of Biotechnology (BT/PR8675/GET/119/1/2015) and Science and Engineering Research Board (EMR/2016/005273), Ministry of Science & Technology, Government of India.

## ASSOCIATED CONTENT

### Supporting Information

Histograms of interaction energies, helical propensities, mean hydrophobic moments and helical wheel projections of BH3-like peptides, vdw and electrostatic components of selected BH3-like peptides, MD snapshots of protein-peptide complexes and data related to cell viability and cell proliferation studies are presented in Fig. S1 to Fig. S19. Details of all the MD simulations is provided in Table S1.

## Abbreviations

Bcl-2: B cell lymphoma 2
BH domain: Bcl-2 homology domain
Bcl-XL: B cell lymphoma – extra large
Mcl-1: Myeloid cell leukaemia 1
Bid: BH3-interacting domain death agonist
Bim: Bcl-2-interacting mediator of cell death
Bad: Bcl-2-associated agonist of cell death
Puma: p53 upregulated modulator of apoptosis
MD: Molecular dynamics

## Notes

### Competing Interest Statement

The authors have declared no competing interest.

