## Supporting Information for "A novel computational method to design BH3-mimetic peptide inhibitors that can bind specifically to Mcl-1 or Bcl-X_L_"

#### Figure S1

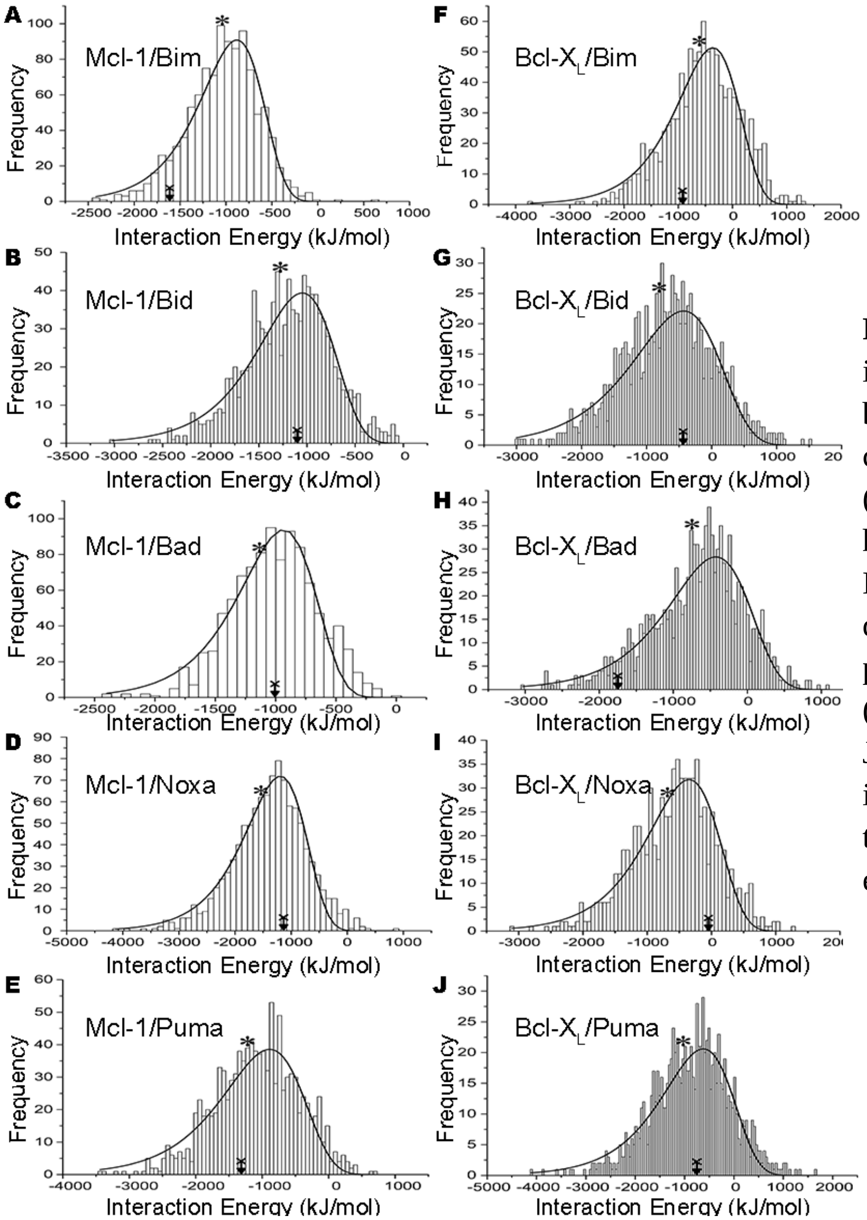


**Figure S1**: Histograms of interaction energies calculated between BH3-like peptides in complex with (A-E) Mcl-1 and (F-J) Bcl-$X_L$. The data shown here correspond to the Set-I of BH3-like peptide sequences derived from the wild-type peptides (A, F) Bim, (B, G) Bid, (C, H) Bad, (D, I) Noxa and (E, J) Puma. The mean value and the interaction energy of the wild-type BH3 peptide are marked in each histogram.

**Figure S2**

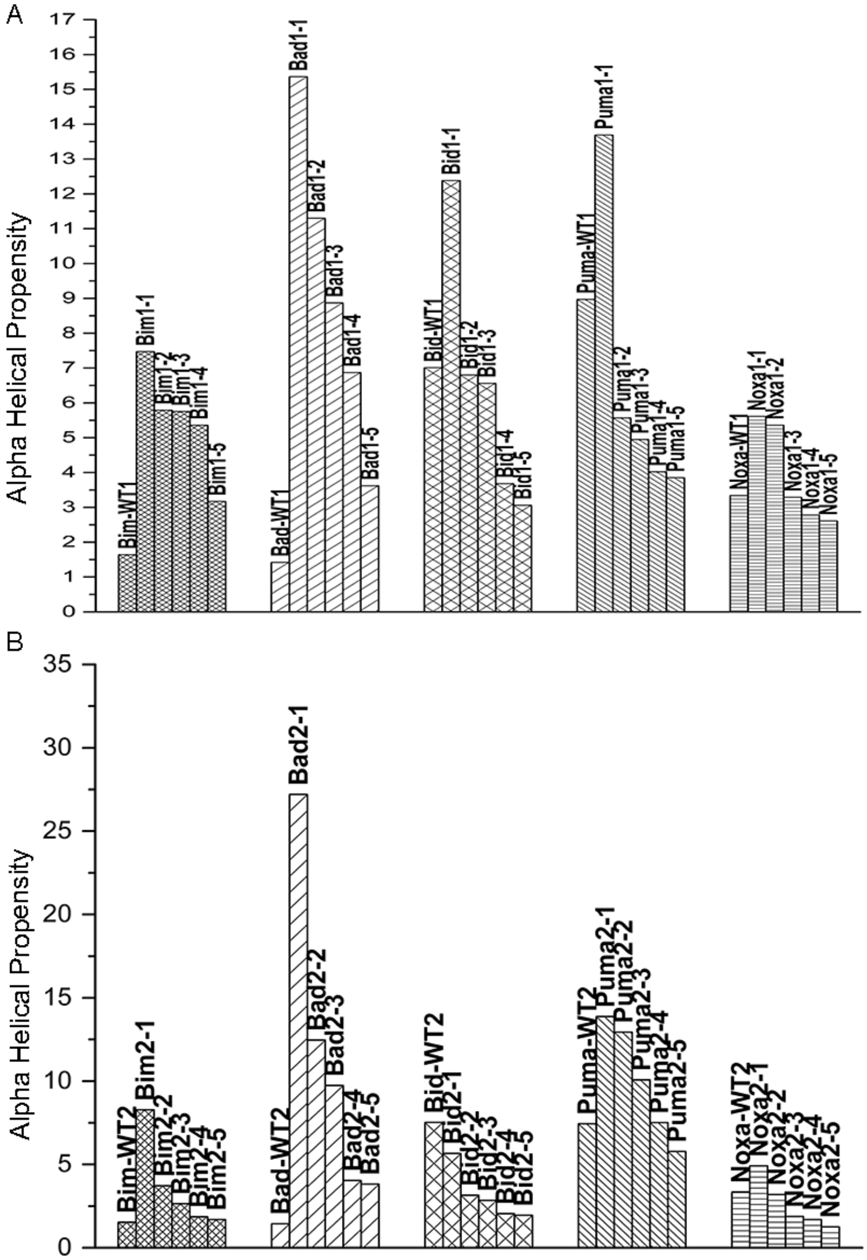


**Figure S2**: Helical propensities of top BH3-like sequences that bind to Mcl-1 with highly favorable interaction energies from Set-I (top) and Set-II (bottom). Helical nature of all BH3-like sequences has been determined using AGADIR web server (http://agadir.crg.es/).

#### Figure S3

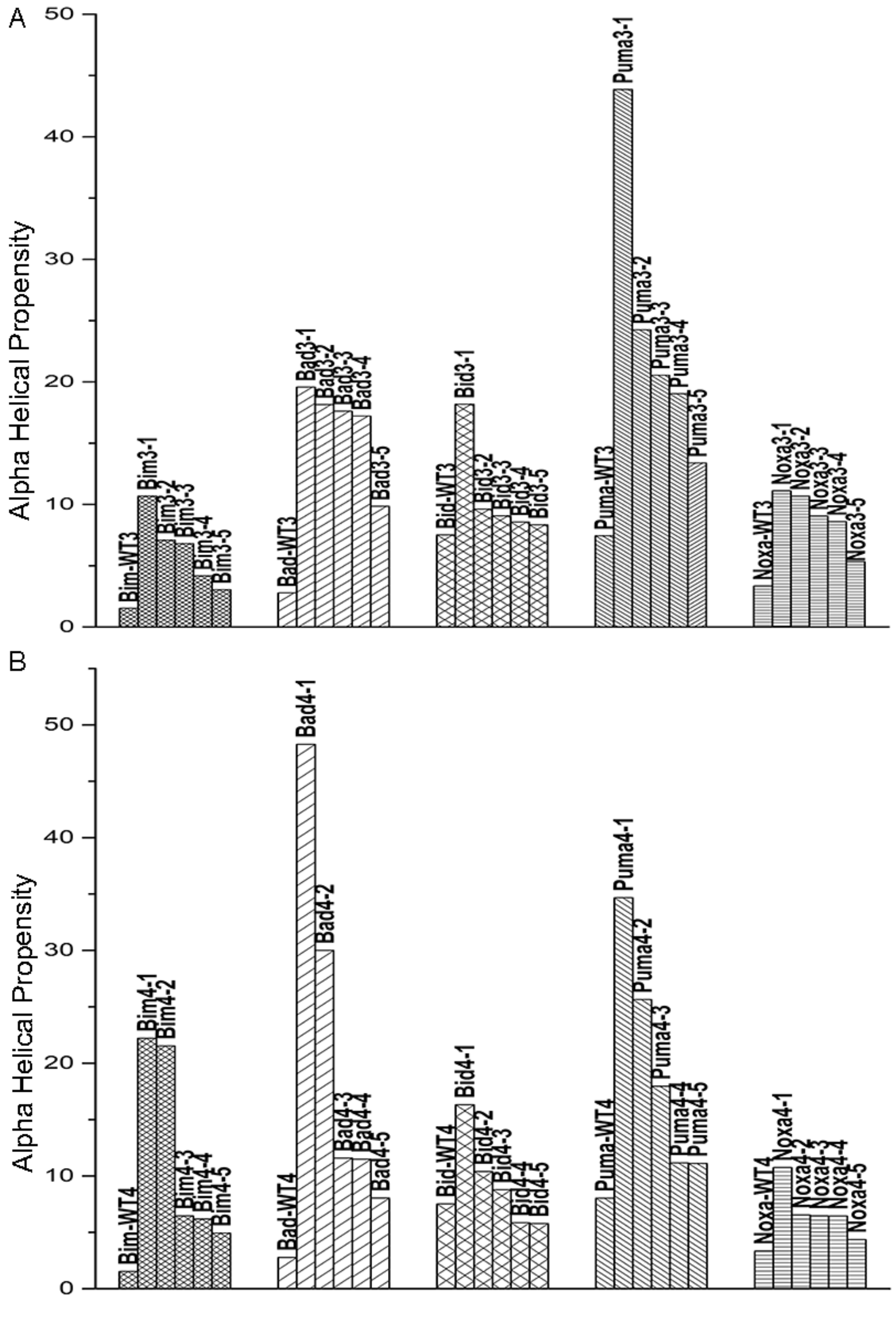


**Figure S3**: Helical propensities of top BH3-like sequences that bind to Bcl-$X_L$ with highly favorable interaction energies from Set-I (top) and Set-II (bottom). Helical nature of all BH3-like sequences has been determined using AGADIR web server (http://agadir.crg.es/).

**Figure S4**

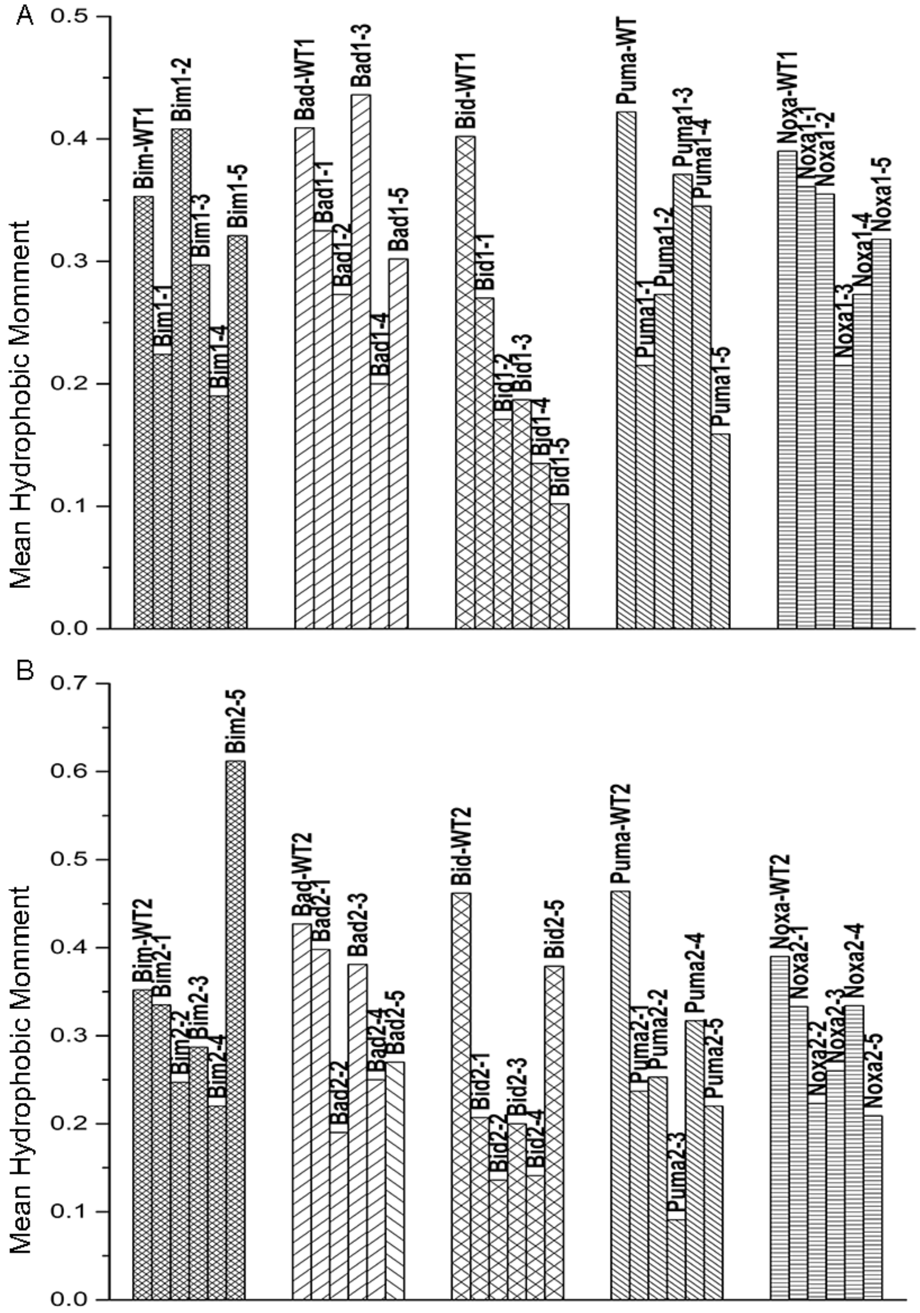


**Figure S4**: Mean helical hydrophobic moment of BH3-like sequences that bind to Mcl-1 with highly favorable interaction energies from Set-I (top) and Set-II (bottom). Mean hydrophobic moment was calculated using the HELIQUEST web server (http://heliquest.ipmc.cnrs.fr/).

### Figure S5

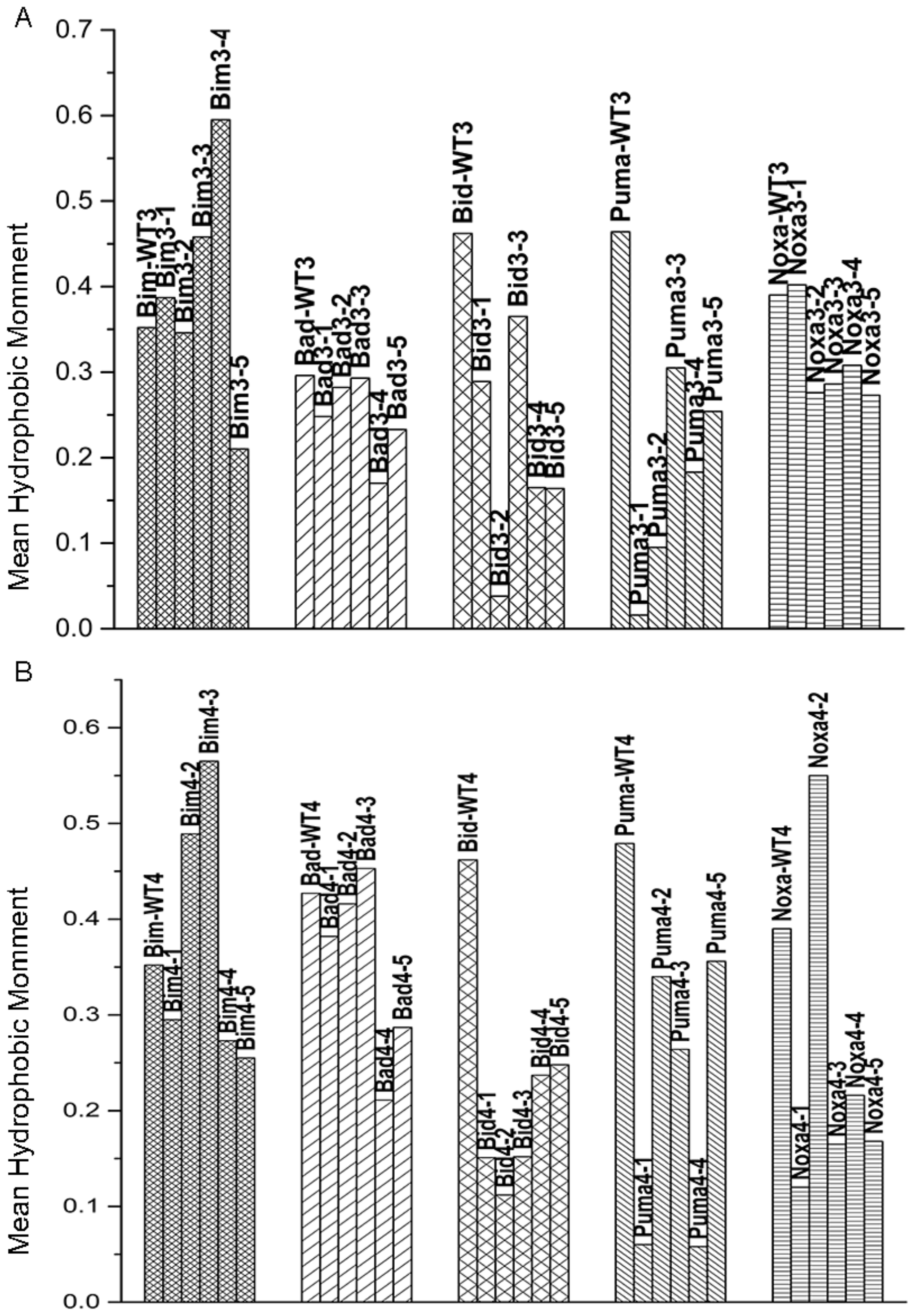


**Figure S5**: Mean helical hydrophobic moment of BH3-like sequences that bind to Bcl-$X_L$ with highly favorable interaction energies from Set-I (top) and Set-II (bottom). Mean hydrophobic moment was calculated using the HELIQUEST web server (http://heliquest.ipmc.cnrs.fr/).

#### Figure S6

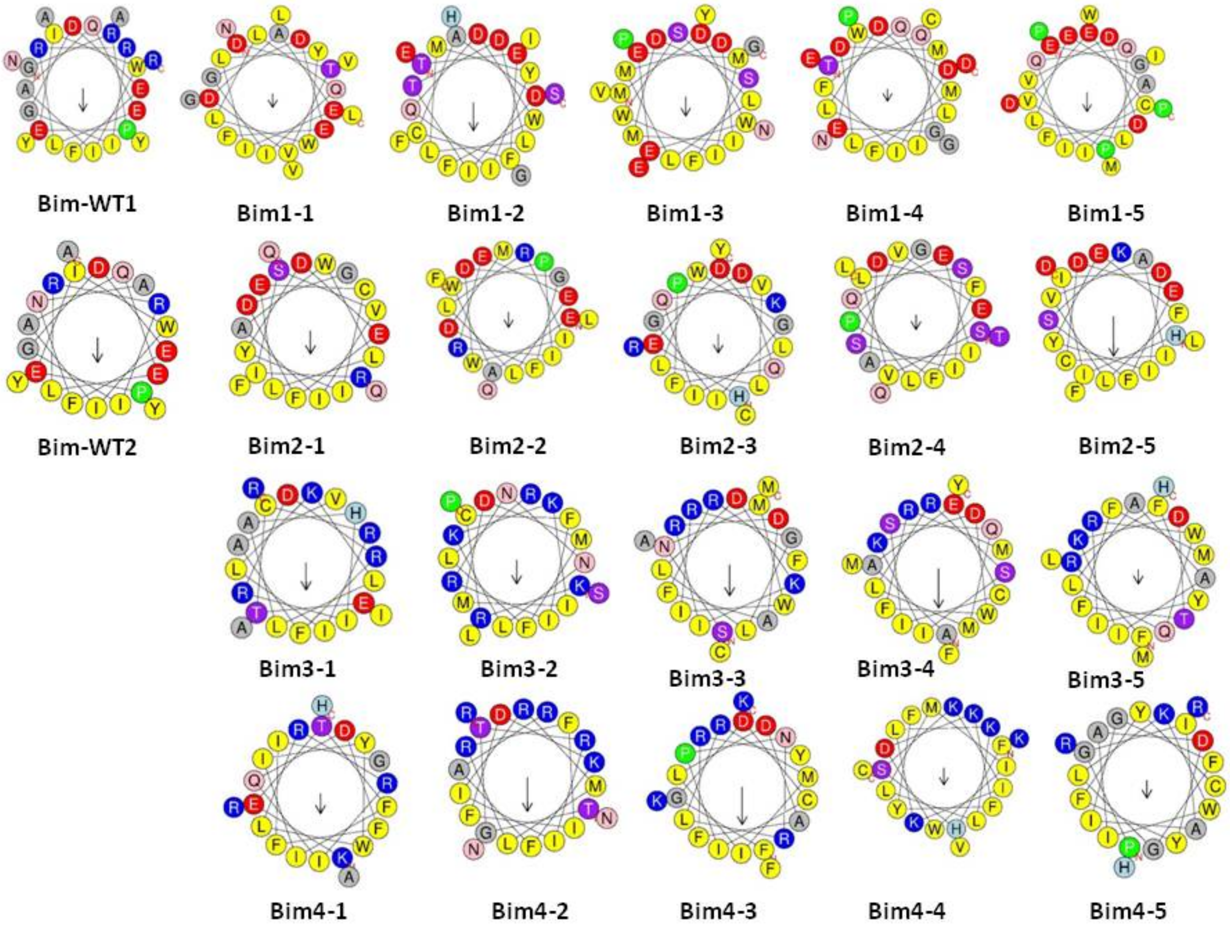


**Figure S6**: Helical wheel projections of Bim wild-type peptides and Bim-based BH3-like sequences.

#### Figure S7

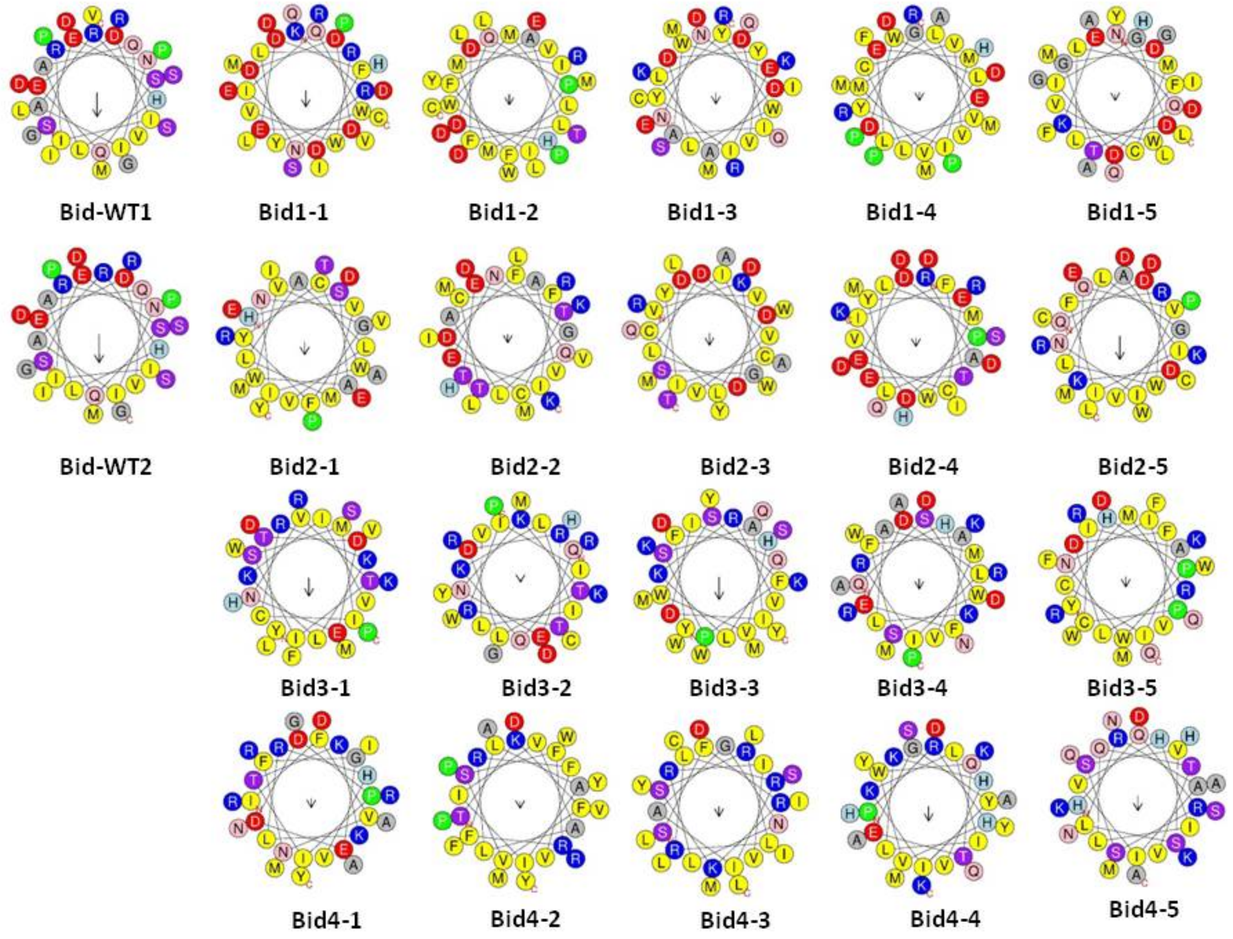


**Figure S7**: Helical wheel projections of Bid wild-type peptides and Bid-based BH3-like sequences.

### Figure S8

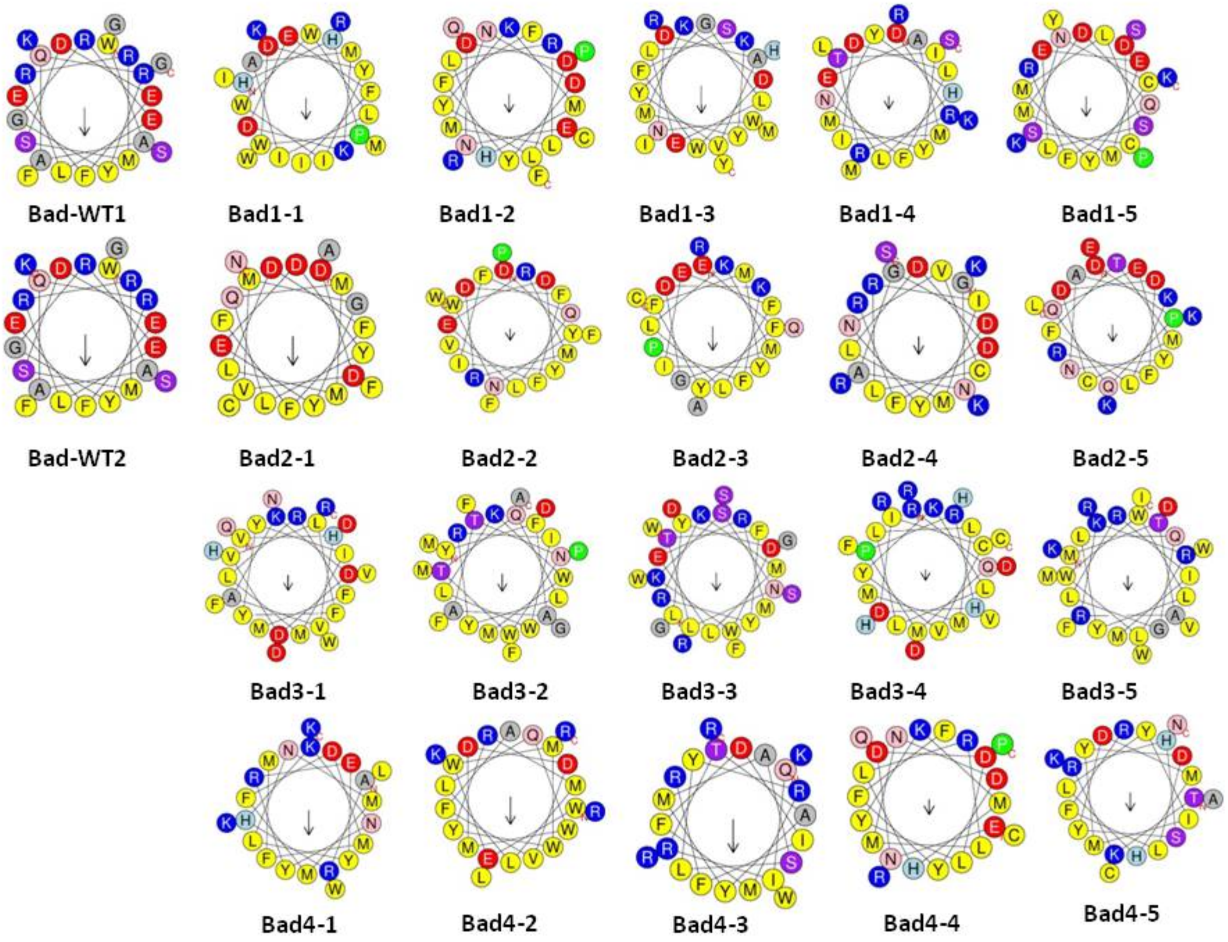


**Figure S8**: Helical wheel projections of Bad wild-type peptides and Bad-based BH3-like sequences.

#### Figure S9

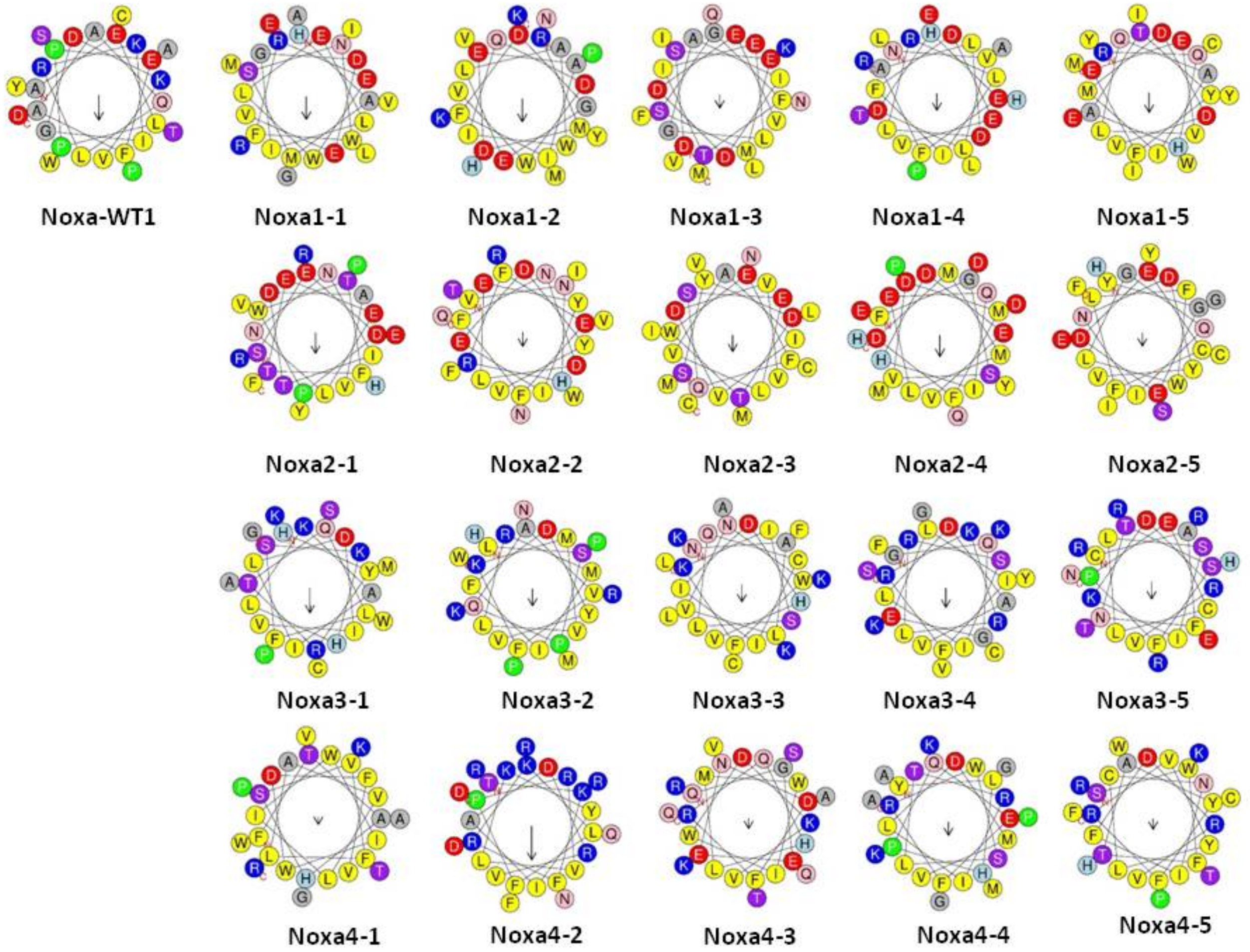


**Figure S9**: Helical wheel projections of Noxa wild-type peptide and Noxa-based BH3-like sequences

#### Figure S10

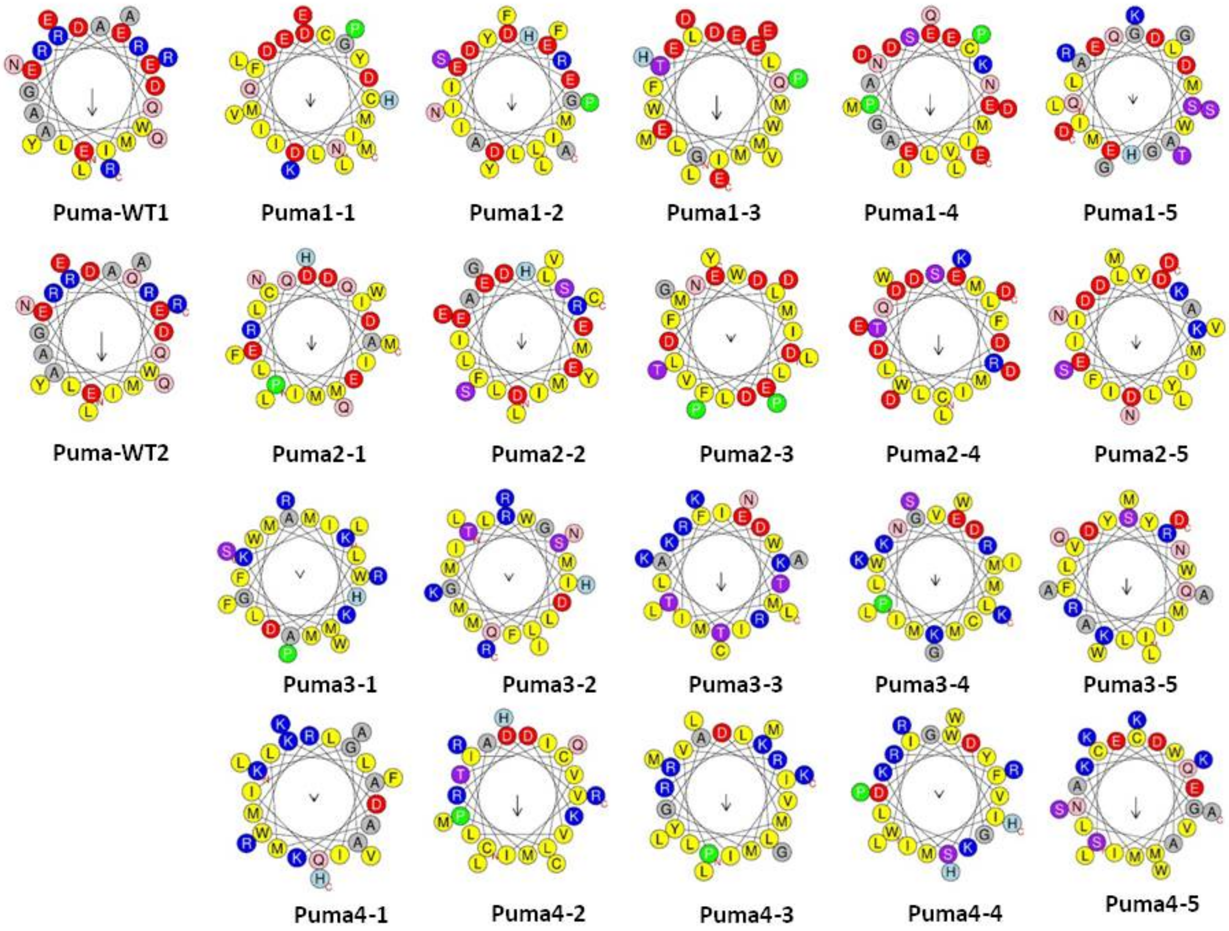


**Figure S10**: Helical wheel projections of Puma wild-type peptides and Puma-based BH3-like sequences

#### Figure S11

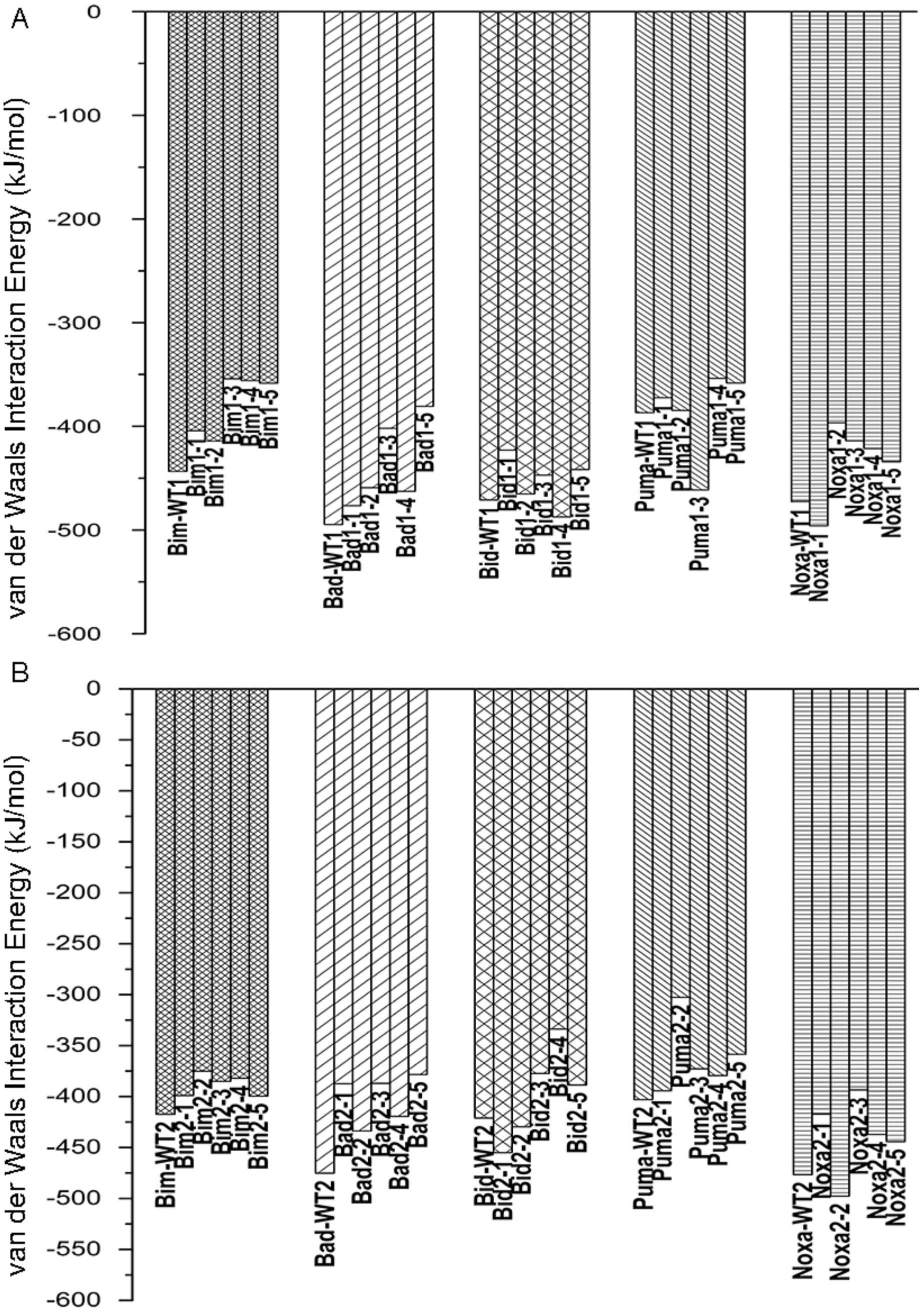


**Figure S11:** van der Waals component of interaction energies between the top BH3-like peptides and the Mcl-1 protein shown for Set-I (top) and Set-II (bottom).

### Figure S12

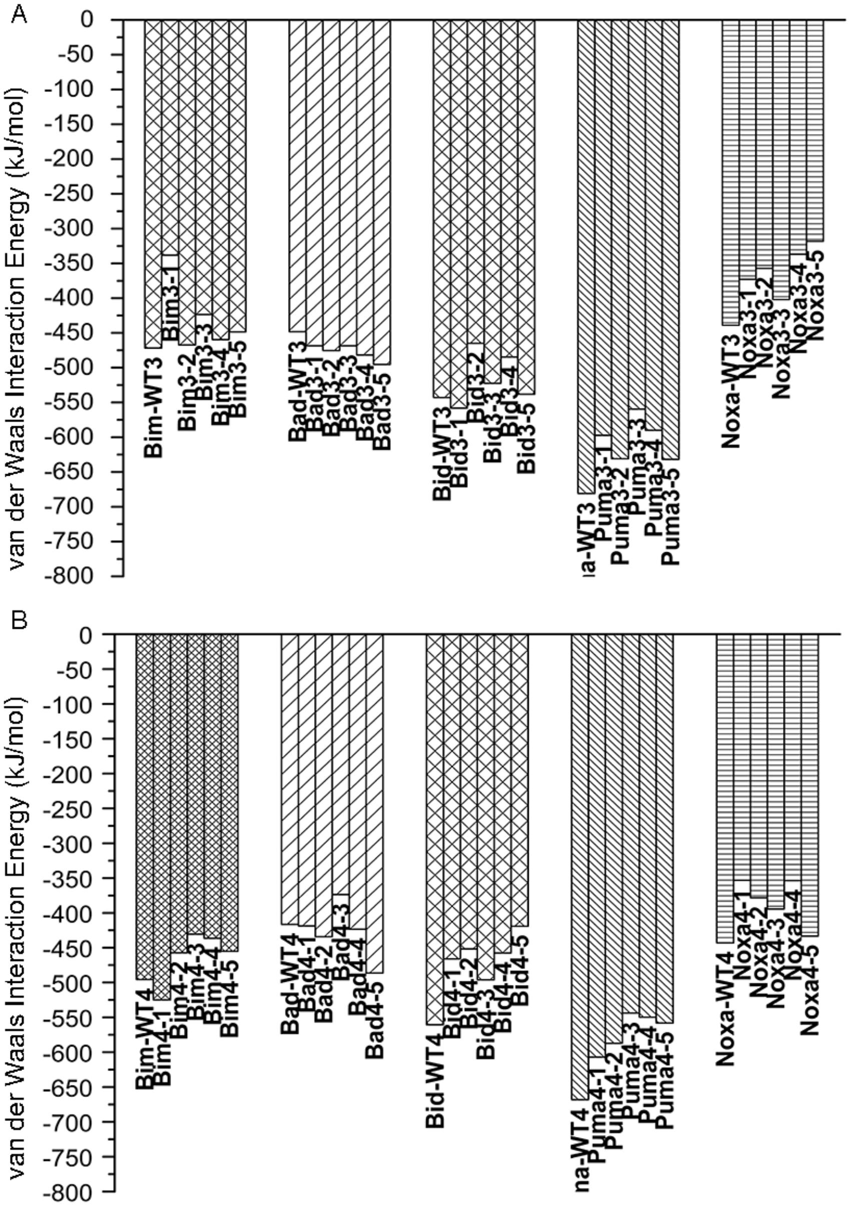


**Figure S12:** van der Waals component of interaction energies between the top BH3-like peptides and the Bcl-$X_L$ protein shown for Set-I (top) and Set-II (bottom).

**Figure S13**

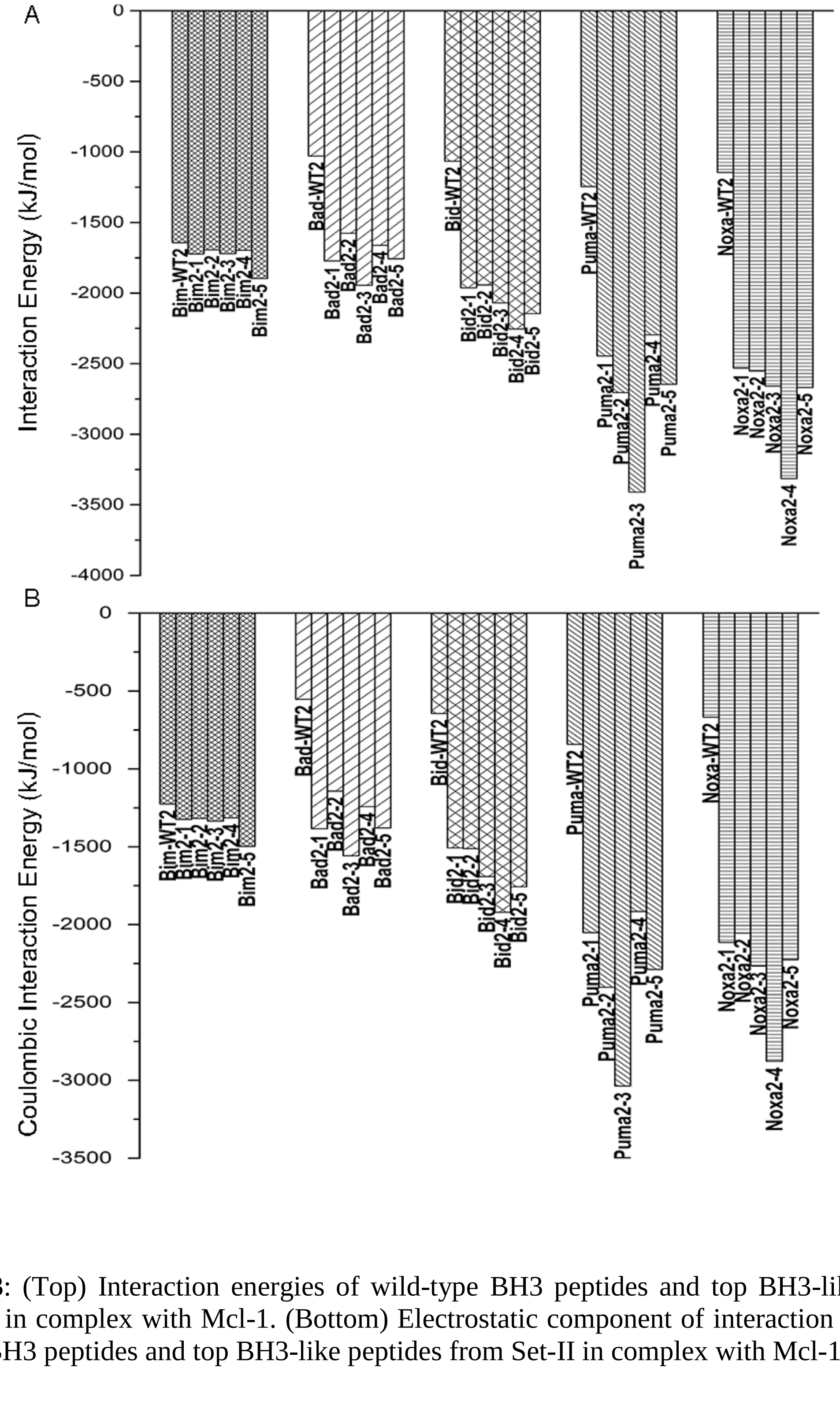


**Figure S13**: (Top) Interaction energies of wild-type BH3 peptides and top BH3-like peptides from Set-II in complex with Mcl-1. (Bottom) Electrostatic component of interaction energies of wild-type BH3 peptides and top BH3-like peptides from Set-II in complex with Mcl-1

#### Figure S14

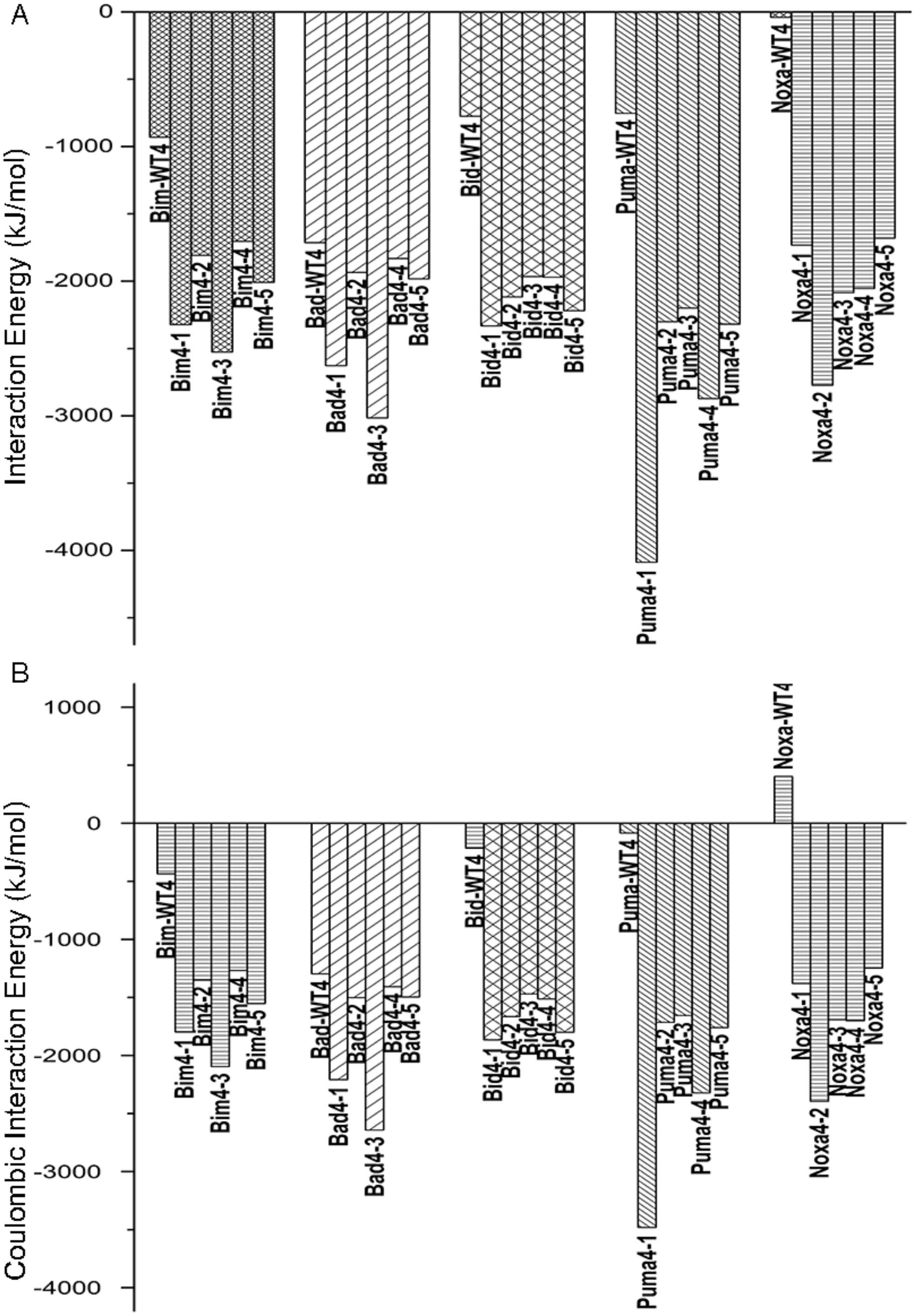


**Figure S14**: (Top) Interaction energies of wild-type BH3 peptides and top BH3-like peptides from Set-II in complex with Bcl-$X_L$. (Bottom) Electrostatic component of interaction energies of wild-type BH3 peptides and top BH3-like peptides from Set-II in complex with Bcl-$X_L$.

#### Figure S15

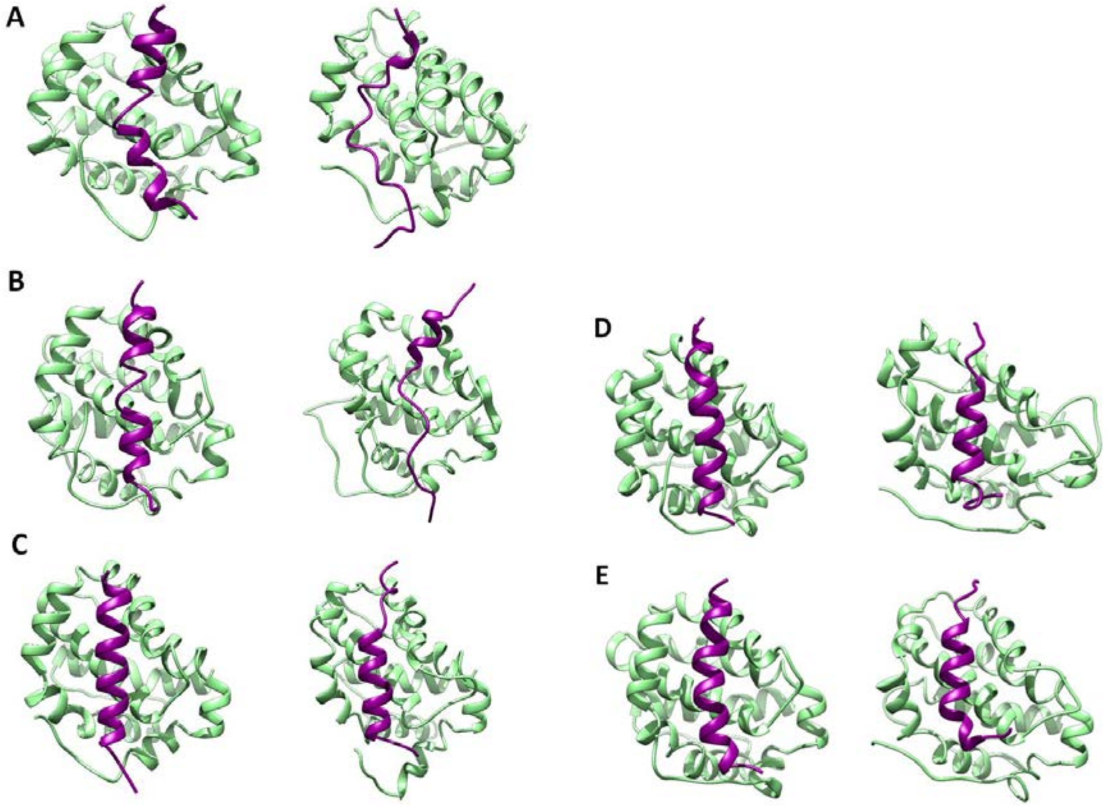


**Figure S15**: Mcl-1 in complex with Bad-WT1 and Bad2-1 peptides. (A-C) Starting structures and the structures saved at the end of 500 ns simulations of Mcl-1 protein in complex with Bad-WT1 are shown for three independent simulations (Mcl-1:Bad-WT1/Sim1, Mcl-1:Bad-WT1/Sim2 and Mcl-1:Bad-WT1/Sim3). (D-E) Starting structures and the structures saved at the end of 500 ns production runs of Mcl-1 protein in complex with Bad2-1 peptide are shown for two independent simulations (Mcl-1:Bad2-1/Sim1, Mcl-1:Bad2-1/Sim2). For summary of all MD simulations, see Table S1.

**Figure S16**

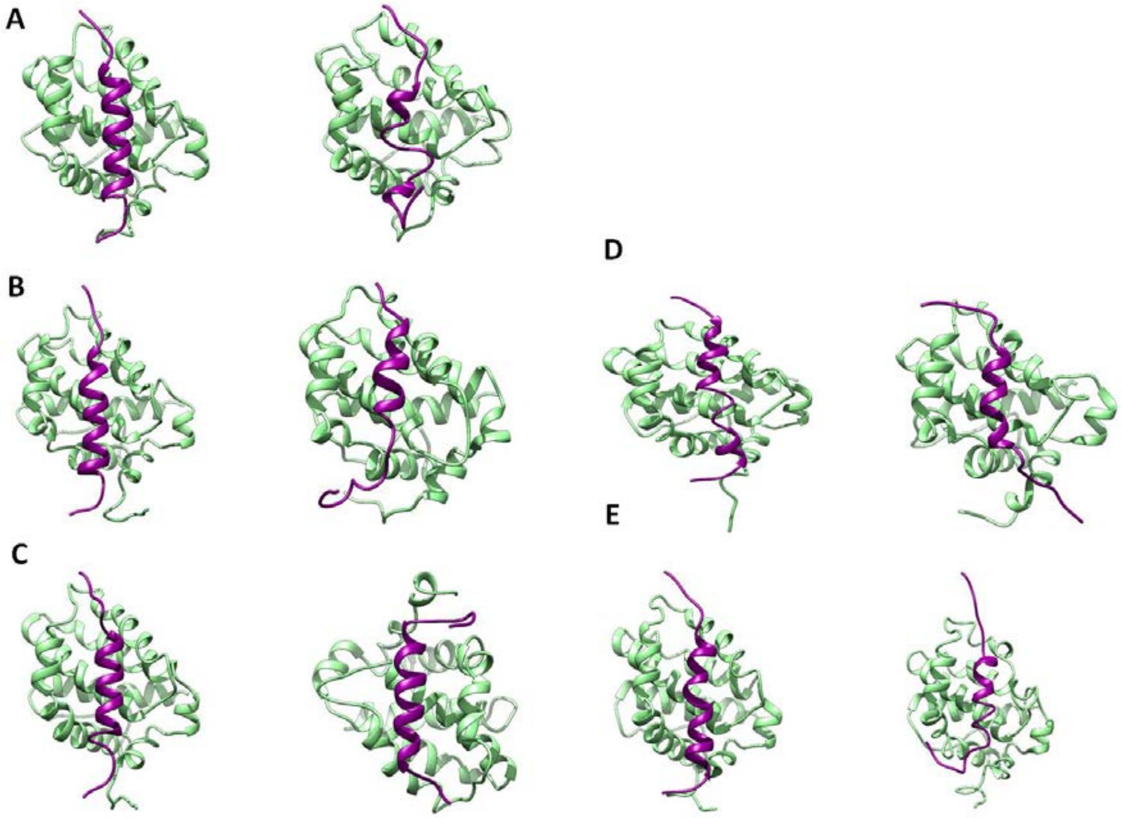


**Figure S16**: Mcl-1 in complex with Noxa-WT1 and Noxa3-1 peptides. (A-C) Starting structures and the structures saved at the end of 500 ns simulations of Mcl-1 protein in complex with Noxa-WT1 are shown for three independent simulations (Mcl-1:Noxa-WT1/Sim1, Mcl-1:Noxa-WT1/Sim2 and Mcl-1:Noxa-WT1/Sim3). (D-E) Starting structures and the structures saved at the end of 500 ns production runs of Mcl-1 protein in complex with Noxa3-1 peptide are shown for two independent simulations (Mcl-1:Noxa3-1/Sim1 and Mcl-1:Noxa3-1/Sim2). For summary of all MD simulations, see Table S1.

#### Figure S17

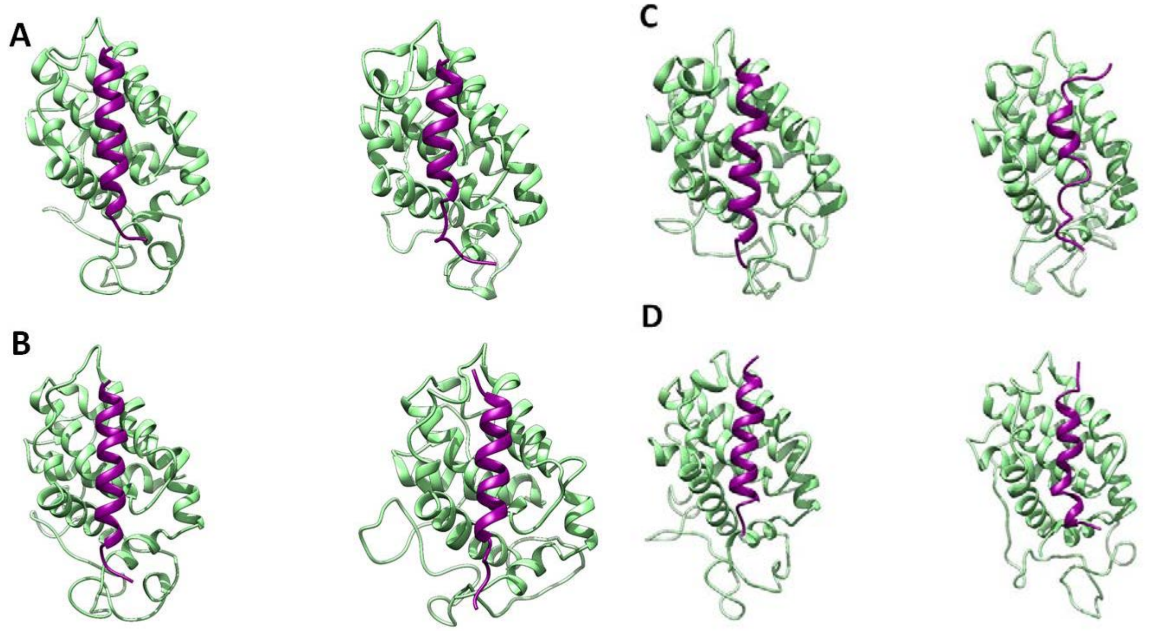


**Figure S17**: Bcl-$X_L$ in complex with Bad-WT1 and Bad2-1 peptides. (A-B) Starting structures and the structures saved at the end of 500 ns simulations of Bcl-$X_L$ protein in complex with Bad-WT1 are shown for two independent simulations (Bcl-$X_L$:BadWT1/Sim1 and Bcl-$X_L$:Bad-WT1/Sim2). (C-D) Starting structures and the structures saved at the end of 500 ns production runs of Bcl-$X_L$ protein in complex with Bad2-1 peptide are shown for two independent simulations (Bcl-$X_L$:Bad2-1/Sim1 and Bcl-$X_L$:Bad2-1/Sim2). For summary of all MD simulations, see Table S1.

#### Figure S18

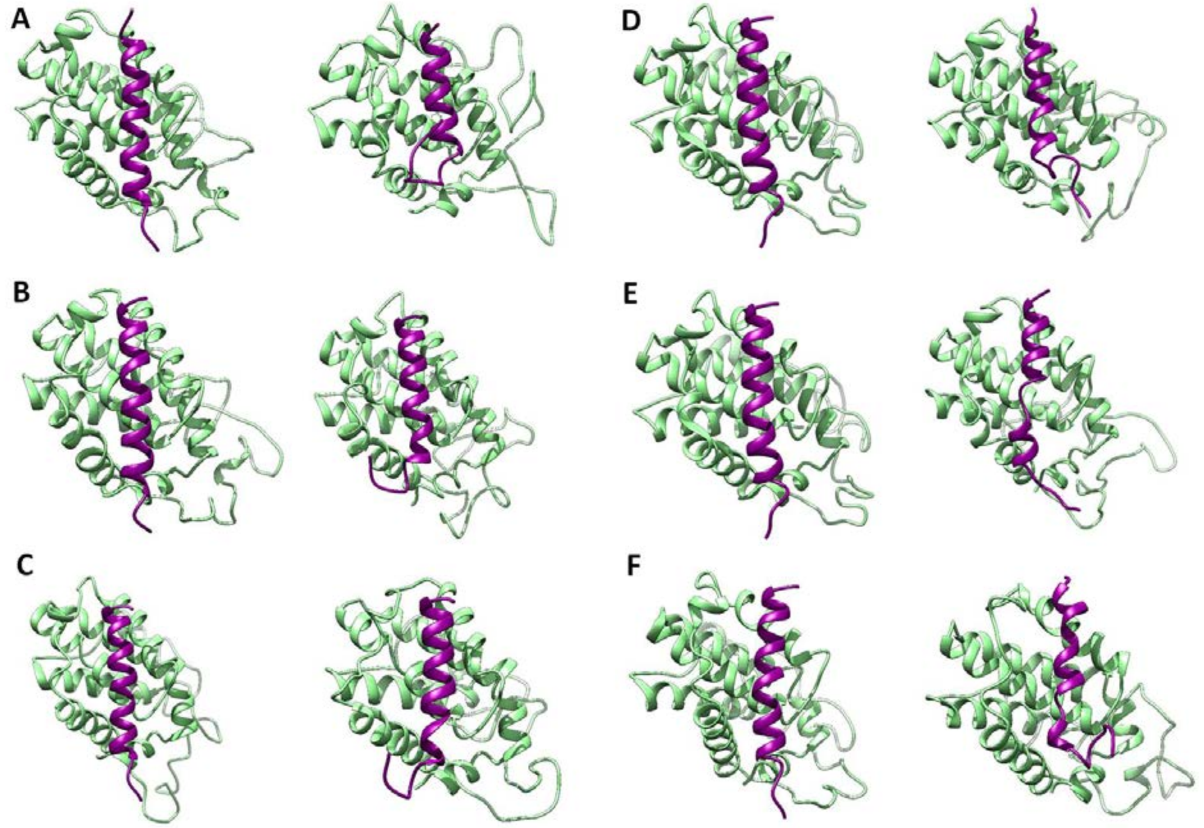


**Figure S18**: Bcl-$X_L$ in complex with Noxa-WT1 and Noxa3-1 peptides. (A-C) Starting structures and the structures saved at the end of 500 ns simulations of Bcl-$X_L$ protein in complex with Noxa-WT1 are shown for three independent simulations (Bcl-$X_L$:NoxaWT1/Sim1, Bcl-$X_L$:Noxa-WT1/Sim2 and Bcl-$X_L$:Noxa-WT1/Sim3). (D-F) Starting structures and the structures saved at the end of 500 ns production runs of Bcl-$X_L$ protein in complex with Noxa3-1 peptide are shown for three independent simulations (Bcl-$X_L$:Noxa3-1/Sim1, Bcl-$X_L$:Noxa-3-1/Sim2 and Bcl-$X_L$:Noxa-3-1/Sim3). For summary of all MD simulations, see Table S1.

**Figure S19**

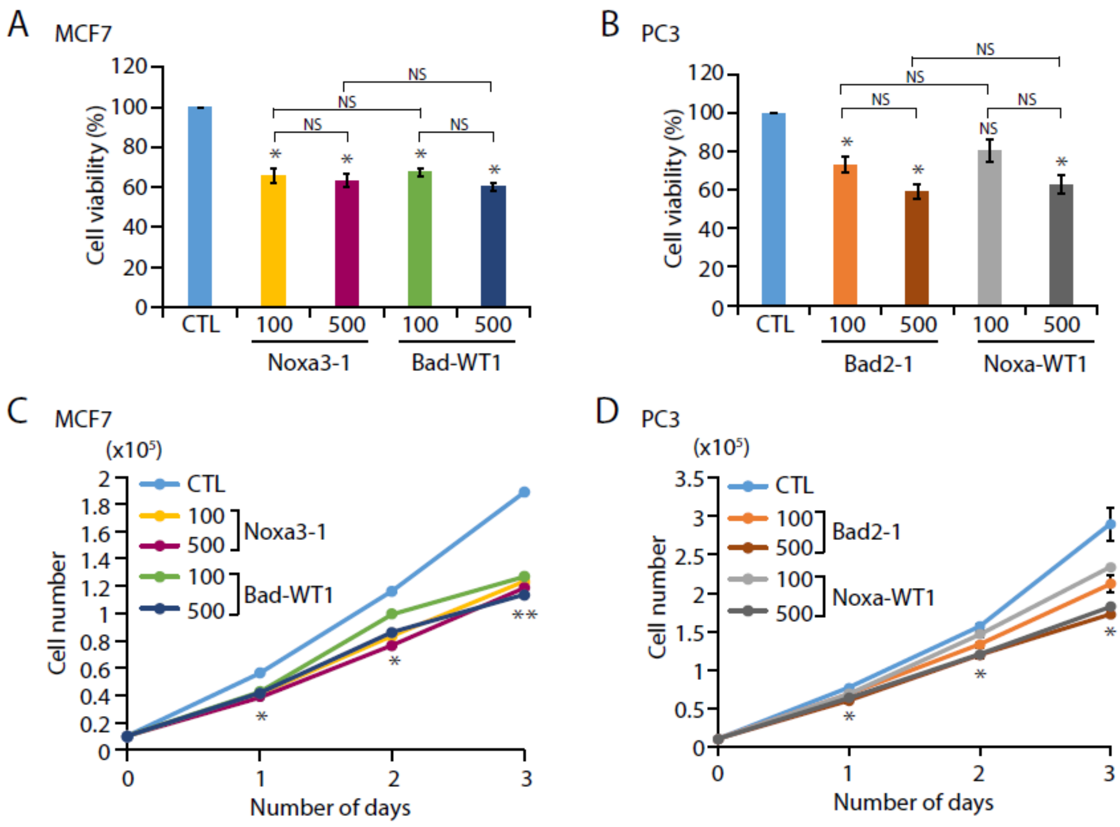


**Figure S19:** Synthetic BH3-like peptides against Mcl-1 and Bcl-$X_L$ reduce cancer cell growth. (**A**) Percent cell viability using MCF7 cells treated with different concentrations (100nM and 500nM) of Noxa3-1 and Bad-WT1 peptides along with control (CTL). (**B**) Percent cell viability using PC3 cells treated with different concentrations (100nM and 500nM) of Bad2-1 and Noxa-WT1 peptides along with CTL. (**C**) Cell proliferation assay using same cells as in (A). (**D**) Cell proliferation assay using same cells as in (B).

**Table S1**: Summary of molecular dynamics simulations of BH3-like and wild-type BH3 peptides in complex with Mcl-1 and Bcl-XL

| S. No. | Protein | Peptide | Simulation | Production run |
|---|---|---|---|---|
| 1. | Mcl-1 | Noxa-WT1 | Mcl-1:Noxa-WT1/Sim1 | 500 ns |
| 2. | Mcl-1 | Noxa-WT1 | Mcl-1:Noxa-WT1/Sim2 | 500 ns |
| 3. | Mcl-1 | Noxa-WT1 | Mcl-1:Noxa-WT1/Sim3 | 500 ns |
| 4. | Mcl-1 | Bad-WT1 | Mcl-1:Bad-WT1/Sim1 | 500 ns |
| 5. | Mcl-1 | Bad-WT1 | Mcl-1:Bad-WT1/Sim2 | 500 ns |
| 6. | Mcl-1 | Bad-WT1 | Mcl-1:Bad-WT1/Sim3 | 500 ns |
| 7. | Mcl-1 | Bad2-1 | Mcl-1:Bad2-1/Sim1 | 500 ns |
| 8. | Mcl-1 | Bad2-1 | Mcl-1:Bad2-1/Sim2 | 500 ns |
| 9. | Mcl-1 | Noxa3-1 | Mcl-1:Noxa3-1/Sim1 | 500 ns |
| 10. | Mcl-1 | Noxa3-1 | Mcl-1:Noxa3-1/Sim2 | 500 ns |
| 11. | Bcl-$X_L$ | Noxa-WT1 | Bcl-$X_L$:Noxa-WT1/Sim1 | 500 ns |
| 12. | Bcl-$X_L$ | Noxa-WT1 | Bcl-$X_L$:Noxa-WT1/Sim2 | 500 ns |
| 13. | Bcl-$X_L$ | Noxa-WT1 | Bcl-$X_L$:Noxa-WT1/Sim3 | 500 ns |
| 14. | Bcl-$X_L$ | Bad-WT1 | Bcl-$X_L$:Bad-WT1/Sim1 | 500 ns |
| 15. | Bcl-$X_L$ | Bad-WT1 | Bcl-$X_L$:Bad-WT1/Sim2 | 500 ns |
| 16. | Bcl-$X_L$ | Bad2-1 | Bcl-$X_L$:Bad2-1/Sim1 | 500 ns |
| 17. | Bcl-$X_L$ | Bad2-1 | Bcl-$X_L$:Bad2-1/Sim2 | 500 ns |
| 18. | Bcl-$X_L$ | Noxa3-1 | Bcl-$X_L$:Noxa3-1/Sim1 | 500 ns |
| 19. | Bcl-$X_L$ | Noxa3-1 | Bcl-$X_L$:Noxa3-1/Sim2 | 500 ns |
| 20. | Bcl-$X_L$ | Noxa3-1 | Bcl-$X_L$:Noxa3-1/Sim3 | 500 ns |
